# The translational role of SOS1 in colorectal cancer

**DOI:** 10.1101/2022.10.14.512156

**Authors:** Diego Alem, Xinrui Yang, Francisca Beato, Bhaswati Sarcar, Alexandra F. Tassielli, Ruifan Dai, Tara L. Hogenson, Margaret A. Park, Kun Jiang, Jianfeng Cai, Yu Yuan, Martin E. Fernandez-Zapico, Aik Choon Tan, Jason B. Fleming, Hao Xie

## Abstract

**Background:** It has been challenging to develop agents directly targeting *KRAS* driver mutations in colorectal cancer (CRC). Recent efforts have focused on developing inhibitors targeting SOS1 as an attractive approach for *KRAS*-mutant cancers. Here, we aimed to study the translational role of SOS1 in CRC using patient-derived organoids (PDOs).

**Method:** In this study, we used CRC PDOs as preclinical models to evaluate their sensitivity to SOS1 inhibitor BI3406 and its cellular effects. We utilized large CRC datasets including GENIE cohort, TCGA PanCancer Atlas, and CPTAC-2 cohort to study the significance of molecular alterations of SOS1 in CRC. To identify potential predictive markers, we performed immunohistochemistry (IHC) on CRC tissue for SOS1/2 protein expression and RNA sequencing to identify discriminative gene sets for sensitivity to SOS1 inhibition. The findings were validated by DepMap data for *SOS1* dependency.

**Result:** CRC PDOs instead of cell lines had differential sensitivities to BI3406. There was a significant correlation between SOS1 and SOS2 mRNA expressions (Spearman’s ρ 0.56, p<0.001). SOS1/2 protein expression by IHC were universal with heterogeneous patterns in cancer cells but only minimal to none in surrounding non-malignant cells. SOS1 protein expression was associated with worse overall survival in patients with *RAS/RAF* mutant CRC (*p*=0.04). We also found that SOS1/SOS2 protein expression ratio > 1 by IHC (*p*=0.03) instead of *KRAS* mutation (*p*=1) was a better predictive marker to BI3406 sensitivity of CRC PDOs. This was concordant with the significant correlation between SOS1/SOS2 protein expression ratio by mass spectrometry and SOS1 dependency score. RNA-seq and gene set enrichment analysis revealed differentially expressed genes and 7 enriched gene sets involving cholesterol homeostasis, epithelial mesenchymal transition, and TNFα/NFκB signaling in BI3406-resistant CRC PDOs. We further discovered that GTP-bound RAS level underwent rebound at 48 hours upon treatment with BI3406 even in BI3406-sensitive PDOs with no change of KRAS effector genes downstream. Cellular adaptation mechanisms to SOS1 inhibition may involve upregulation of SOS1/2 mRNA and SOS1 protein expressions, which may be overcome by SOS1 knockdown/degradation or synergistic effect of BI3406 with trametinib.

**Conclusion:** In summary, CRC PDOs could serve as better models for translational study of SOS1 in CRC. High SOS1 protein expression was a worse prognostic factor in CRC. High SOS1/SOS2 protein expression ratio predicted sensitivity to SOS1 inhibition and dependency. Our preclinical findings supported further clinical development of SOS1-targeting agents in CRC.

## Introduction

Approximately half of the metastatic colorectal cancers (CRC) among other cancers carry driver mutations in the *RAS* family genes. They are associated with poor responses to standard chemotherapy and serve as negative predictive marker to anti-EGFR blockade.^1^ As a result, patients with *RAS*-mutant CRC have worse outcome and urgently need novel targeted therapy options. For these reasons, there have been significant efforts and advances recently in the development of novel therapeutics either directly inhibiting mutant KRAS or targeting functionally important *RTK/RAS* effector pathways, namely MAPK and PI3K.^2^ However, similar to other targeted agents in clinical practice, set aside issues with primary resistance, development of acquired resistance is inevitable in majority of the cases and toxicities of combination therapies may become prohibitory to tolerate.^3^ These necessitate the development of novel strategies to overcome these clinical dilemmas.

SOS1 constitutes part of the signaling protein complexes downstream of RTKs to regulate RTK-dependent *RAS* activation as in CRC and EGFR-mutant non-small cell lung cancer (NSCLC). SOS1 can not only promote the production of active GTP-bound RAS at the catalytic site as a guanine nucleotide exchange factor (GEF), but also enhance its own GEF function in an allosteric fashion.^4^ Targeting SOS1 may have advantages over other indirect approaches to *RAS* signaling suppression given its direct protein-protein interaction with RAS.^3^ It may also be less toxic due to functional compensation from its paralog SOS2 and lack of requirement in normal cells in contrast to SHP2.^5^ In addition to small molecule SOS1 agonists developed by Fesik and colleagues,^6–11^ a few of the SOS1 inhibitors have also been discovered including BAY293, BI3406, MRTX0902 and others.^12–16^ BAY293 was able to disrupt SOS1-KRAS interaction and demonstrated cellular activity in various cancer cell lines *in vitro*.^12^ BI3406 was the first potent SOS1 inhibitor with single digit nanomolar binding affinity and *in vivo* activities.^13^ Its related compound BI1701963 is currently under a phase I clinical trial alone or in combination with trametinib in advanced solid tumors.^17^ The availability of these agents provides the opportunity to specifically target part of the *RTK/RAS* signaling proximal to *RAS*, which allows new possibility of combination therapies.

However, most of these SOS1-targeting agents were discovered in cancer cell lines and cell line-derived xenograft models without specific patient context. Preclinical findings on SOS1 inhibition were mainly from NSCLC where SOS1 inhibitor not only had single agent activity but also synergy with EGFR-TKIs and KRAS G12C inhibitors in *EGFR*-mutant or *KRAS G12C* NSCLC cell lines, respectively.^12,18,19^ In the latter case, SOS1 inhibitor can enhance KRAS G12C inhibitor binding to GDP-bound KRAS, decrease KRAS G12C activation, and thus suppress oncogenic signaling.^16^ As to predictive markers, previous studies showed that SOS1 inhibitor as a single agent is active in cell lines, especially NSCLC cell lines with *EGFR* and *KRAS G12/13* mutations instead of *KRAS Q61* and *BRAF* type I/II mutations.^12,13,19^ But these findings were without further characterizations in patient-relevant cancer models. In contrast to NSCLC, *RTK/RAS* signaling in CRC is very different given its intrinsic upregulation of RTK ligands. SOS1 inhibitor as a single agent was less effective in inhibiting *RAS*-driven CRC cell growth, which required combination with a MEK inhibitor for enhanced *in vivo* activity in CRC.^13^

Hence, to define the translational role of SOS1 and SOS1 inhibitors, we evaluated their biological activities and associated predictive markers to sensitivity and resistance in CRC patient-derived organoids (PDOs) as these models closely represent the disease.^20^ PDOs are 3D tumor models resembling key molecular and biological features of the original tumor with superior performance at predicting response to therapy in CRC compared to cell lines or xenograft models.^21–23^ In this study, we demonstrated the potential values of early adoption of PDOs in the discovery of biomarkers and cellular effects associated with SOS1 inhibitors in CRC.

## Materials and Methods

### Patients

Patients were enrolled under prospective protocols approved by Moffitt Cancer Center (MCC) Institutional Review Board including MCC20584 “Generation and *Ex vivo* Testing of Immune-Based Cell Therapy from Gastrointestinal Malignancies” and MCC19990 “Biological determinants of colorectal cancer outcomes in Latinos of diverse ancestral origins”. These protocols allowed the collection of surgically resected tumor specimens including CRC at MCC. Tumor specimens were from either primary or metastatic CRC including but not limited to liver and peritoneal metastases as part of routine clinical care. The use of reagents derived from these tumor specimens was approved under protocol MCC20880 “Preclinical Testing of SOS1 Inhibition and Degradation in *RAS*-Mutant Colorectal Cancer”. Tumor specimens collected were de-identified and assigned a lab number. The type and site of tumor specimen, patient’s demographic and clinical information, treatment history, previous tumor genetic information, and organoid initiation date were collected when available. Patient-derived xenografts (MCC IACUC protocol: 8797R) were used to generate and biobank tumor specimens. The tumor samples were subsequently used to generate additional CRC patient-derived xenografts (PDXs) and PDOs.

### Cell line culture

CRC cell lines HCT116, SW620, SW1417, and C2BBe1 were purchased from the American Type Culture Collection (ATCC) and cultured in modified McCoys’ 5A medium (Gibco, 16600082). MCC20584-T023 was a patient-derived colon cancer cell line from colon cancer ovarian metastasis. It was cultured in RPMI 1640 medium (ATCC, 30-2001) and authenticated by STR profiling using established algorithms.^24^ The culture media were supplemented with 10% FBS (ATCC, 30-2020) and 1% penicillin-streptomycin (ATCC, 30-2300). All cell lines were regularly checked for mycoplasma contamination. They were incubated under a 5% (v/v) CO_2_ atmosphere at 37°C.

### Cell line drug sensitivity assay

A defined-number of cells were plated in a 96-well plate in culture media. Cells were then treated with drugs at various concentrations for 72 hours. Drugs used included BI3406 (Selleckchem, S8916), buparlisib (MedChemExpress, HY-70063), defactinib (MedChemExpress, HY-12289) and trametinib (Selleckchem, S2673). Celltiter-Glo 2.0 reagent (Promega, G9241) was diluted into PBS per manufacture’s protocol and added to the wells. The plate was covered with foil and incubated at room temperature for 10 minutes. The plate was read on an Envision multi-well plate reader (PerkinElmer). After normalization of readings to 0.2% DMSO-treated cells, dose-response curves were generated with IC_50_ values calculated using GraphPad Prism 8.4.3.

### Cell line spheroid drug sensitivity assay

A 96-well plate was treated with 50µL 1.2% agarose, then was added 30,000 cells/well. The plate was incubated under a 5% (v/v) CO_2_ atmosphere at 37°C overnight. Drug solutions at various concentrations in the media were added and incubated for 96 hours. PrestoBlue cell viability reagent (Thermo Fisher Scientific, A13261) was added to the wells. The plate was incubated at culture conditions for 2 hours prior to the read of fluorescence at excitation/emission 560/590nm. Dose-response curves with IC_50_ values were obtained in a similar fashion to the cell line drug sensitivity assay.

### Synergy assessment for drug combinations

Two drugs at various concentrations were combined to treat cells in a similar fashion to the cell line drug sensitivity assay. The relative inhibition values were used as the input for SynergyFinder (https://synergyfinder.fimm.fi)^25^ to calculate the synergy score and most synergistic area score of drug combination using Bliss method.^26,27^ Dose-response matrix (inhibition) and the distribution of Bliss synergy scores were plotted.

### Establishment of CRC PDXs

NOD/SCID and nude mice (female, aged 6 week) were purchased from Jackson Laboratories. Fresh tumor samples were cut into fragments of about 2–3 mm,^28^ briefly soaked in Matrigel, and then implanted in the subcutaneous space of the mice.^29,30^ Tumors were measured weekly. Tumor volumes were calculated using the formula length × width_2_ × 0.52. The tumors were harvested when they reached 0.75-1.5 cm in diameter and labeled F1 to F4 to indicate different generational passages in animals.

To estimate and compared CRC PDX tumor growth, area under the tumor growth curve up to time t (aAUC) was calculated using R script with access provided by.^31^

### Patient-derived organoid culture

CRC organoids were generated and expanded using a protocol similar to previously published protocols for CRC with modifications.^23,32^ Briefly, the tumor specimen was minced into approximately 1 mm^3^ small fragments using sterile scalpels in fresh wash media. Tissue fragments were placed in warmed digestion media and incubated for 30-45 minutes on a shaker with agitation at 37°C and 600 rpm to allow tissue to dissociate into single cells. Larger tissue fragments were allowed to settle under normal gravity. The supernatant was transferred out followed by an addition of 3mL wash media with 10% FBS. Cells were filtered through a 100μM mesh filter and a 40μM filter to remove mouse fibroblasts. Cell pellets suspended in 300μL of ice-cold growth factor reduced Matrigel (Corning, 354230), then plated into 50µL domes in a 24-well pre-warmed culture plate, which was allowed to solidify for 15 minutes at 37°C. When the Matrigel domes solidified, 500µL of pre-warmed complete growth media was added and incubated in 5% CO_2_ atmosphere at 37°C. The growth of the organoids was monitored with fresh growth media change every 2 days. Organoid propagation was performed in a sequence of gentle mechanical digestion followed by enzymatic digestion. The cell pellet was resuspended in wash media with 0.1% BSA, followed by embedment into Matrigel domes. Wash media: advanced DMEM/F12 (Gibco, 12634010), 1M Hepes (Gibco, 15630080), 100X glutamine (Gibco, 25030149), primocin (Invivogen, ANTPM1). Digestion media: wash media/10% FBS, collagenase and dispase (Sigma, 10269638001). Growth media: wash media without FBS, 1X wnt3a/R-spondin/Noggin condition medium (L-WRN),^33^ 1X B27 supplement (Gibco, 12587-010), 10mM nicotinamide (Sigma Aldrich, N3376), 1.25mM N-acetylcysteine (Sigma Aldrich, A9165), 100μg/mL primocin (Invivogen, ANTPM1), 100ng/mL recombinant mouse Noggin (abcam, ab281818), 50ng/mL hEGF (R&D Systems, 236EG200), 100ng/mL human FGF (R&D Systems, 233-FB), 10nM human gastrin I (R&D Systems, 3006/1), 500nM A83-01 (Selleckchem, S7692), 10.5μM Y-267632 (Selleckchem, S1049), and 1μM PGE2 (R&D Systems, 2296/10).

### Organoid drug sensitivity assay

Organoids were harvested with organoid cell recovery solution (Corning, 354253) and pipetted gently to dissolve Matrigel. After incubation for 15 minutes, the cells were collected and washed with 0.1% BSA/wash media, followed by resuspension in a mixture of 90% complete growth media and 10% Matrigel. Cells were seeded in triplicates into a 96-well plate previously prepared with solidified 30μL of 50% Matrigel and 50% complete growth media in each well.

Once the organoids were visible after 3-7 days, drugs were added and cultured for 72 hours. Chemiluminescence was read at 360/460nm on an Envision multi-well plate reader (PerkinElmer) after addition of CellTiter-Glo 3D (Promega, G9681). After normalization to 0.2% DMSO-treated cells, dose-response curves were generated with IC_50_ values calculated using GraphPad Prism 8.4.3.

### Immunoblot

Cells were plated in a 100mm dish for incubation overnight before harvest for immunoblot. Cell confluency was checked prior to proceeding with the treatment. Cells were then lysed in 1x radioimmunoprecipitation assay (RIPA) lysis buffer (Thermo Scientific, 89900) and Halt™ Protease and Phosphatase Inhibitor Cocktail (Thermo Scientific, 78429) while maintained on ice. After centrifugation at 12,000 rpm for 15 minutes at 4°C, the protein concentration was determined using the BCA protein assay kit (Thermo Scientific, 23227). Approximately 40μg protein samples were loaded into each well and separated by SDS-PAGE gel. After protein transfer onto nitrocellulose membranes, 5% skim milk in TBST was used to block the membranes for 1 hour at room temperature prior to incubation with primary antibodies overnight at 4°C. At the completion, a horseradish peroxidase–conjugated secondary antibody was applied at room temperature for 1 hour. The bands were visualized using Western Lightning Plus-ECL (PerkinElmer). Primary antibodies used in this study included GAPDH (Cell Signaling, 2118), pERK (Cell Signaling, 4376), ERK (Cell Signaling, 9102), cleaved PARP (Cell Signaling, 5625), beta-actin (abcam, ab8224), and SOS1 (Cell Signaling, 5890).

To prepare siRNA-treated samples for immunoblot, cells were plated with media for 24 hours, then treated with 12μL lipofectamine RNAiMAX (Invitrogen: 13778150) in 200μL Opti-MEM (Gibco: 31985070) and 4μL siRNA (10μM) in 200μL Opti-MEM. The mixture was incubated at room temperature for 5 minutes. After 48 hours, the cells were harvested and processed for immunoblot. siRNAs used included SOS1 (ambion, s13286, 4390824), GAPDH (ambion, s5572, 4390824), and negative control (Invitrogen: AM4613).

### Immunohistochemistry of tumor tissue for SOS1 and SOS2

Tumor tissue blocks were paraffin-embedded and subjected to serial sections for hematoxylin and eosin staining and immunohistochemistry (IHC) following a common procedure at the histology lab. The slides/sections were cut at 4-micron thickness using a Leica RM2245 microtome. Charged slides were used.

SOS1 IHC: Slides were stained using a Ventana Discovery XT automated system (Ventana Medical Systems) as per manufacturer’s protocol with proprietary reagents. Briefly, slides were deparaffinized on the automated system with Discovery Wash Solution. Heat-induced antigen retrieval method was used in Cell Conditioning 1 Standard. The rabbit primary antibody that reacts to SOS1, (Abcam, ab140621) was used at a 1:50 concentration in Dako antibody diluent (Carpenteria, CA) and incubated for 2 hours. The Ventana OmniMap anti-rabbit secondary antibody was used for 16 minutes. The detection system used was the Ventana ChromoMap kit and slides were then counterstained with hematoxylin. Slides were then dehydrated and coverslipped as per normal laboratory protocol. Positive control was performed in SOS1-expressing breast carcinoma. Negative control was performed using substitution of isotype-specific immunoglobulins at the same protein concentration as the primary antibody.

SOS2 IHC: Slides were prepared in a similar fashion as described for SOS1 IHC. A blocking agent was used, Neuvision Block (Ventana) for 16 minutes. The rabbit primary antibody that reacts to SOS2 (Invitrogen, PA5-61488) was used at a 1:100 concentration in Dako antibody diluent (Carpenteria, CA) and incubated 32 minutes. The Ventana OmniMap anti-rabbit secondary antibody was used for 16 minutes. The detection system used was the Ventana ChromoMap kit and slides were then counterstained with hematoxylin. Slides were then dehydrated and coverslipped as per normal laboratory protocol. Positive control was performed in SOS2-expressing colon cancer. Negative control was performed using substitution of isotype-specific immunoglobulins at the same protein concentration as the primary antibody.

Slides were reviewed by a gastrointestinal pathologist (K. J.) who adopted a semiquantitative scoring system of IHC results as previously described.^34^ Briefly, percentage of positively stained cells was assigned a score as 0 = 0%, 1 = <30%, 2 = 30-60%, 3 = >60%. Intensity of IHC stain was assigned a score as 0 = no reaction, 1 = weak, 2 = mild, 3 = strong. Multiplication of the two scores provided the final score.

### Ras GTPase level in organoids

Ras GTPase Chemi ELISA Kit (Active Motif, 52097) was used to measure Ras GTPase level in organoids. The assay was performed according to the manufacture’s manual. Briefly, organoids were treated with 1µM BI3406 and harvested at different time points (0, 6, 24 and 48 hours) for the preparation of whole-cell extract. GST-Raf-RBD diluted in complete lysis binding buffer was added to each well. The plate was incubated for 1 hour at 4°C. Test extract diluted in complete lysis/binding buffer along with positive control extract (EGF treated HeLa) and complete lysis/binding buffer were added to the corresponding wells. Diluted Ras antibody, specific for human HRAS and KRAS, was added to the wells. The plate was incubated for 1 hour at room temperature. Diluted HRP antibody was added to all wells. The plate was then incubated for 1 hour at room temperature, followed by addition of chemiluminescent working solution. The chemiluminescence was read by a luminometer.

### Next generation sequencing using QiaSeq cancer panel

Qiagen AllPrep kit (Qiagen, 80284) was used for dual RNA/DNA extraction. DNA extracted from cells was quantified with the Qubit Fluorometer (ThermoFisher Scientific, Waltham, MA) and screened for quality on the Agilent TapeStation 4200 (Agilent Technologies, Santa Clara, CA). In order to assess the presence of somatic mutations, targeted sequencing was performed using the 275-gene Qiagen QIAseq Comprehensive Cancer Panel (Qiagen, Germantown, MD). Briefly, 40 ng of DNA was enzymatically fragmented followed by followed by end-repair, A-addition, and adaptor ligation. Target enrichment with locus-specific primers and library amplification was performed, and the final libraries were screened on an Agilent TapeStation (Agilent Technologies, Inc., Santa Clara, CA) and quantitated by qPCR with the Kapa Library Quantification Kit (Roche Diagnostics, U.S., Indianapolis, IN). The indexed samples were sequenced on the MiSeq sequencer (Illumina, Inc. San Diego, CA) with 150-base paired-end reads in order to generate >1400X average target coverage after PCR duplicate removal with the use of unique molecular indexes (UMIs). Data analysis including alignment and variant calling were performed using the QiaSeq data analysis pipeline.

### RNA-sequencing and data analysis

RNA extracted from cells was quantitated with the Qubit Fluorometer (ThermoFisher Scientific, Waltham, MA) and screened for quality on the Agilent TapeStation 4200 (Agilent Technologies, Santa Clara, CA). The samples were then processed for RNA-sequencing using the NuGEN Universal RNA-Seq Library Preparation Kit with NuQuant (Tecan Genomics, Redwood City, CA). Briefly, 100ng of RNA was used to generate cDNA and a strand-specific library following the manufacturer’s protocol. Quality control steps were performed, including TapeStation size assessment and quantification using the Kapa Library Quantification Kit (Roche, Wilmington, MA). The final libraries were normalized, denatured, and sequenced on the Illumina NextSeq 2000 sequencer with the P3-200 cycle reagent kit in order to generate approximately 50M million 100-base read pairs per sample (Illumina, Inc., San Diego, CA).

Read adapters were detected using BBMerge (v37.88)^35^ and subsequently removed with cutadapt (v1.8.1)^36^. Processed raw reads were then aligned to human genome HG38 using STAR (v2.5.3a).^37^ Gene expression was evaluated as read count at gene level with RSEM (v1.3.0)^38^ and Ensembl Gencode gene model v32. Gene expression data were then normalized and differential expression between experimental groups were evaluated using DEseq2.^39^ Pathway enrichment were analyzed with gene set enrichment analysis (GSEA).^40^

### mRNA measurement and quantification

Cells were treated with drugs for indicated duration of time prior to harvest for RNA extraction and purification using Qiagen RNeasy mini kit (Qiagen, 74104). The RNA samples were quantified by optical density 260/280, 260/230 readings using a spectrophotometer (NanoDrop, Thermo Fisher Scientific, Waltham, MA, USA). We used the QuantiTect reverse transcription kit (Qiagen, 205311) for cDNA synthesis and gDNA removal in accordance with the manufacturer’s instructions. Quantitative RT-PCR was performed using the MX3000p Real-Time PCR System (Agilent Technologies, Santa Clara, CA, USA) to determine the mRNA expression levels of SOS1, SOS2, and GAPDH. Quantitative RT-PCR for each gene was performed using the TaqMan method with 5μL Taqman fast advanced master mix, 3.5μL nuclease free water, 1μL cDNA template with 0.5μL premade primer sets (*SOS1*: 4331182 Hs00893128_m1, *SOS2*: 4331182 Hs01127273_m1, GAPDH: 4331182 Hs03929097_g1; Thermo Scientific). This was followed by Amplitaq activation at 50°C for 2 minutes, 95°C for 2 minutes, and then 40 cycles at 95°C for 1 second for denaturing and at 60°C for 20 seconds for annealing and extension. We calculated ΔCt, defined as the difference between the crossover threshold (Ct) of the target gene and the Ct average of *GAPDH* for each sample.

### Database used for analyses

We analyzed relevant data for the following publicly available database in this study. They are GENIE cohort v11.0, https://genie.cbioportal.org/.^41,42^ DFCI CRC cohort, https://www.cbioportal.org/study/summary?id=coadread_dfci_2016.^43^ TCGA PanCancer Atlas, https://www.cbioportal.org/study/summary?id=coadread_tcga_pan_can_atlas_2018.^44^ CPTAC-2 Prospective, https://www.cbioportal.org/study/summary?id=coad_cptac_2019.^45^ DepMap, https://depmap.org/portal/.^46^

### Statistical analysis

Statistical analysis for this study was descriptive in nature without sample size or power calculation. Categorical data were summarized as frequency counts and percentages and compared with χ_2_ or Fisher’s exact test. Continuous data were summarized as means, standard deviations, standard errors, medians, and ranges and compared with independent t test or Mann-Whitney U test. For variables with more than two categories, they were compared with one-way ANOVA or Kruskal-Wallis test. Correlation between variables were assessed with Spearman’s rho. All statistical tests were 2-sided. Overall survival (OS) was calculated from the date of tumor resection to the date of death. Surviving patients were censored at the date of last follow-up. Time-to-event data were summarized using Kaplan-Meier method and compared with log-rank tests. Statistical analyses were performed using either GraphPad Prism 8.4.3 or R version 4.2.0.

## Results

### Preclinical models to study SOS1 in CRC

To evaluate the translational role of SOS1 in CRC, we first sought to identify preclinical models that have differential sensitivities to SOS1 inhibitor BI3406. Different from previous studies on testing pan-cancer cell lines, we treated multiple CRC cell lines including a primary CRC cell line MCC20584-T023 derived from colon cancer ovarian metastasis of a patient. As shown in **Figure 1A**, if we define BI3406 sensitivity as <10µM, almost all CRC cell lines were resistant to BI3406 in both 2D and spheroid cultures. The two cell lines, MCC20584-T023 and SW1417, with the lowest IC_50_, 9.85µM and 19.7µM, respectively, had wild-type *KRAS* **(Figure 1B)**.

**Figure 1.**
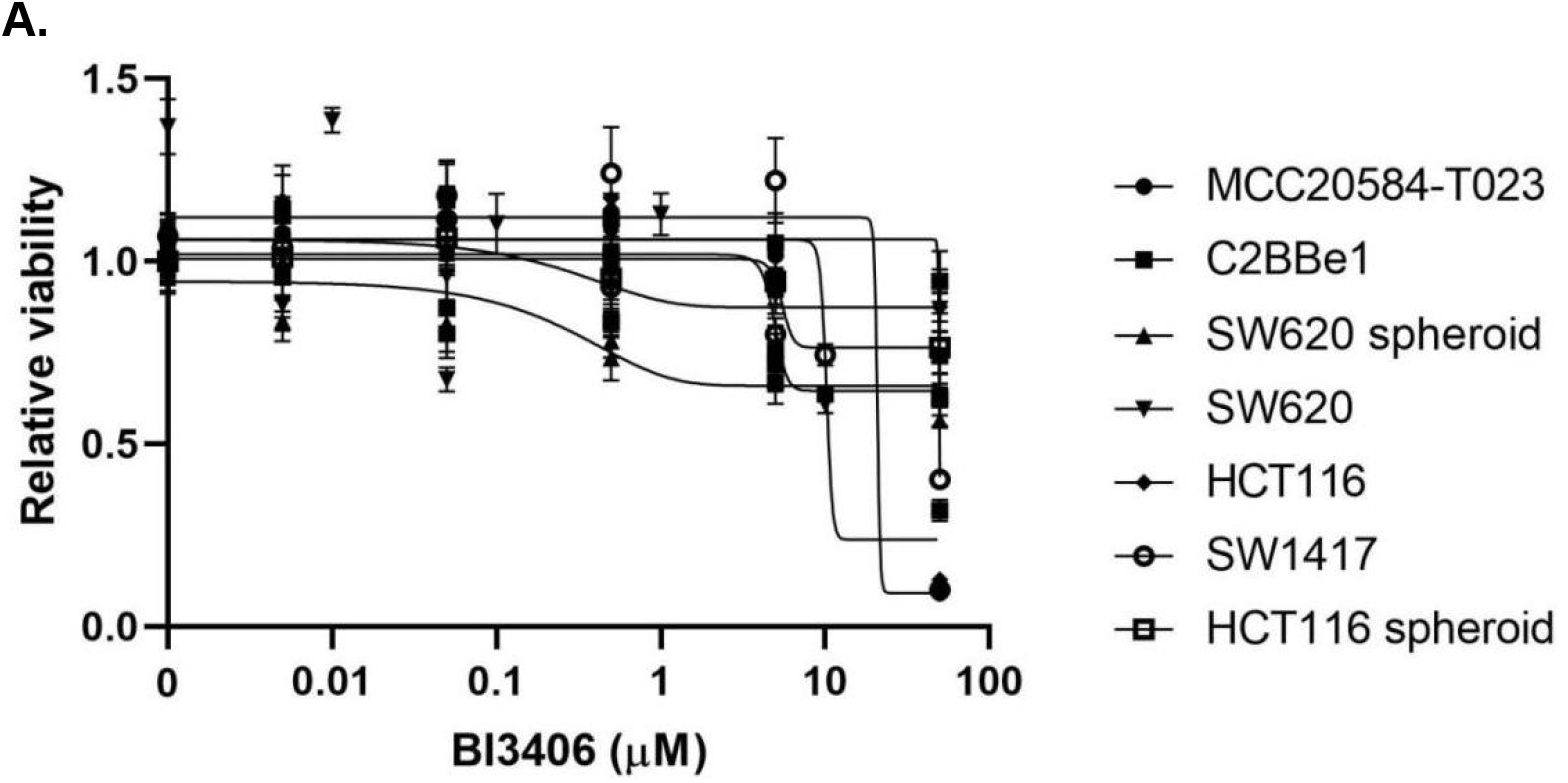
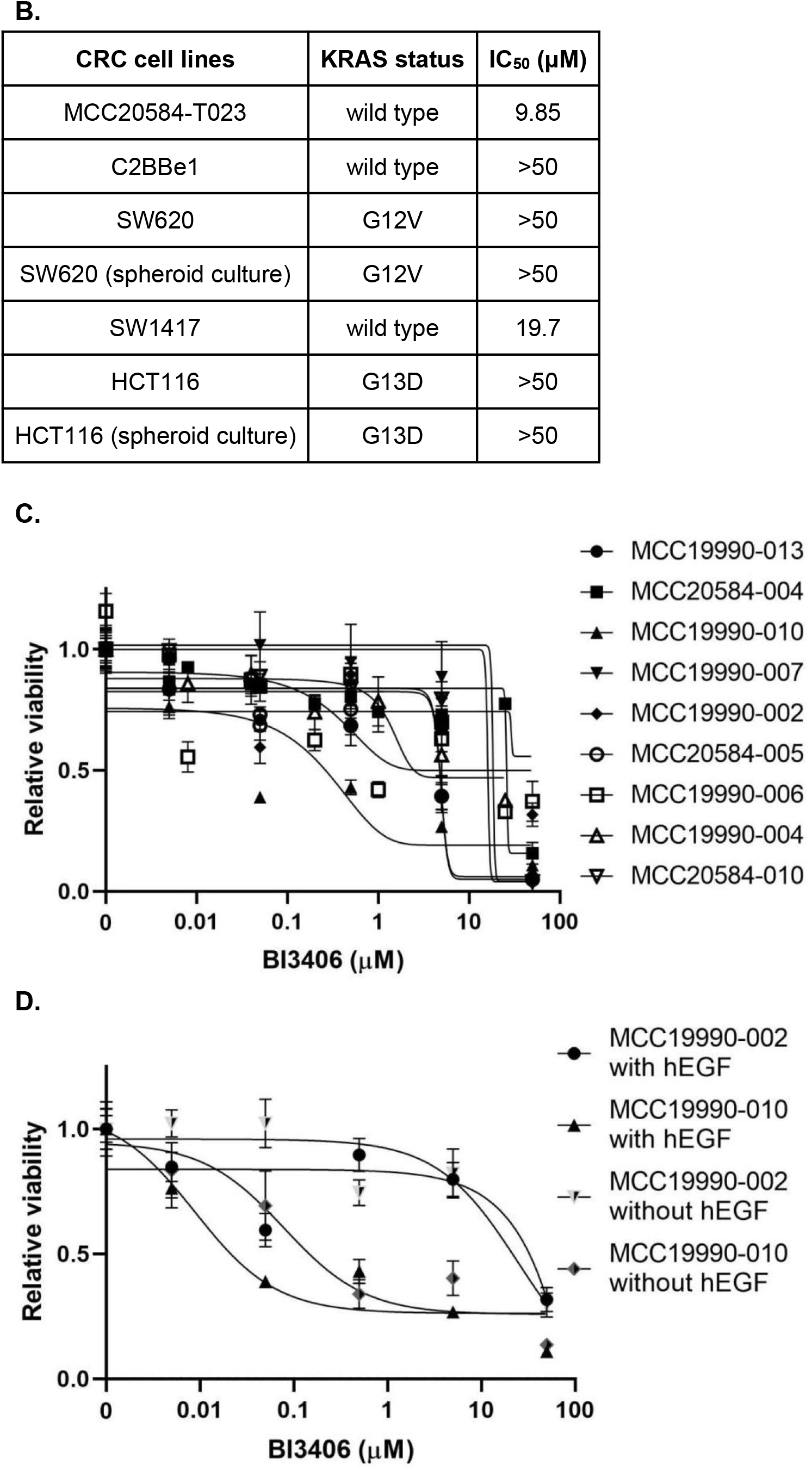
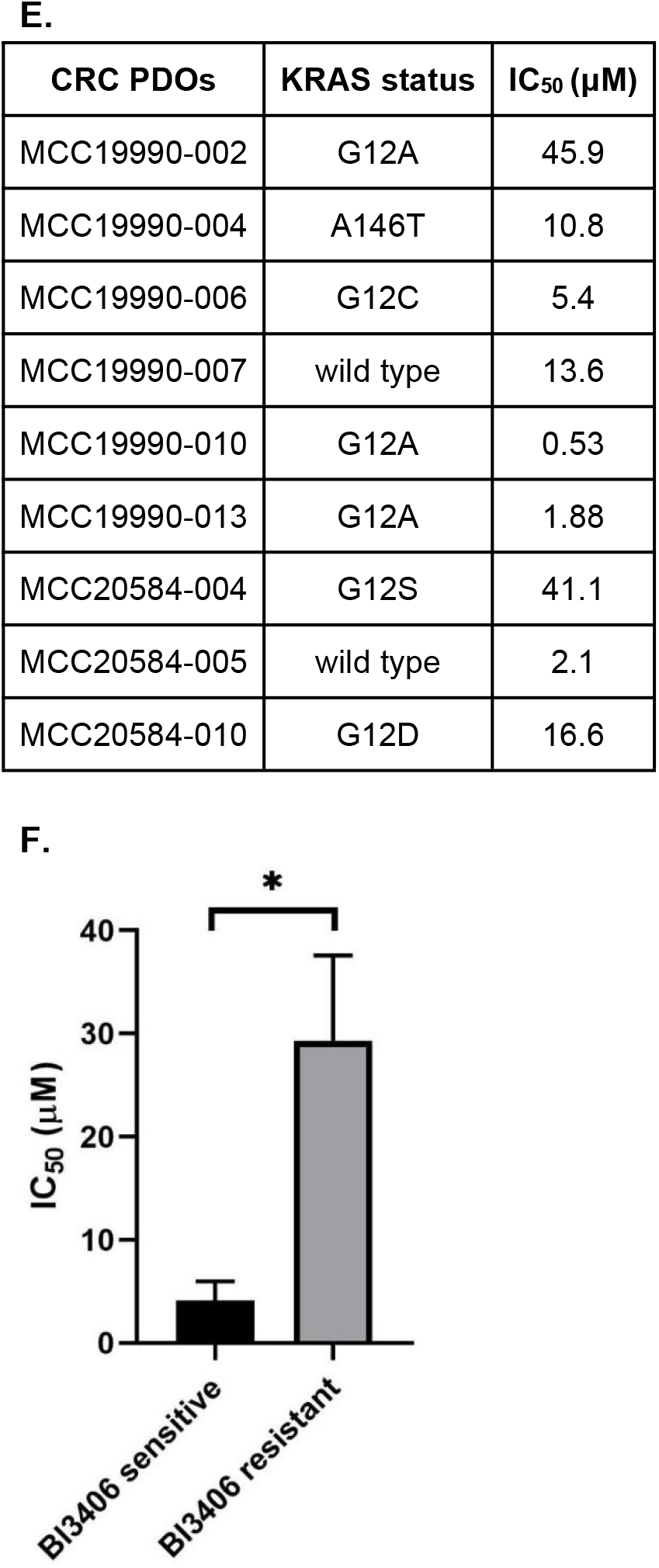
Sensitivity profiles of CRC models to SOS1 inhibitor BI3406. **(A)** Dose-response curves of CRC cell viability to SOS1 inhibitor BI3406. **(B)** IC_50_ values of BI3406 and *KRAS* mutation status in CRC cell lines. **(C)** Dose-response curves of CRC PDOs to BI3406. **(D)** Dose-response curves of selected sensitive and resistant CRC PDOs to BI3406 in the presence or absence of human epidermal growth factor in the culture media. **(E)** IC_50_ values of BI3406 and *KRAS* mutation status in CRC PDOs. **(F)** Comparison of IC_50_ values of BI3406 in SOS1 inhibition sensitive and resistant CRC PDOs. *p*=0.016.

We then assessed the sensitivity of our CRC PDOs to BI3406 and observed a wider distribution of differential sensitivities as evident by the dose-response curves shown in **Figure 1C**. Given the presence of hEGF in organoid culture media, we further looked into the effect of hEGF on SOS1 inhibitor sensitivity in a sensitive CRC PDO MCC19990-010 and a resistant CRC PDO MCC19990-002 **(Figure 1D)**. We found no significant effect of hEGF on SOS1 inhibitor sensitivity. The range of IC_50_ values to BI3406 was between 0.53µM and 45.9µM among 9 CRC PDOs with wild type and various *KRAS* mutations **(Figure 1E)**. These CRC PDOs were derived from surgically resected tumor samples of patients with distinct age, race, gender, tumor location, tumor stage, and microsatellite/mismatch repair protein status **(Supplemental Table 1)**. As shown in **Figure 1F**, the IC_50_ values of BI3406 in BI3406-sensitive CRC PDOs are significantly lower than those in BI3406-resistant CRC PDOs (*p*=0.016).

### Significance of molecular alterations of SOS1 in CRC

In the GENIE cohort (n=5790), the prevalence of SOS1 mutations across different cancer types is generally very low with 2.9% in CRC **(Figure 2A)**. It is not statistically different across different tumor stages (*p*=0.2), primary versus metastatic sites (*p*=0.07), or left versus right sided CRC (*p*=0.3) **(Supplemental Figure 1A-C)**. The prevalence of SOS1 mutations is much lower than that in common clinically relevant genes such as *KRAS, BRAF*, and *HER2*. In addition, SOS1 has co-occurrence with these genes (*q*<0.001) **(Supplemental Figure 1D)** in addition to its co-occurrence with its paralog SOS2 (*q*=0.03) **(Supplemental Figure 1E)**. As to the function of SOS1 mutations, distribution of *SOS1* alterations in CRC of the GENIE cohort was visualized and only 1 of 185 (0.5%) *SOS1* alterations was determined by OncoKB and hotspots as a putative driver of CRC **(Figure 2B)**.

**Figure 2.**
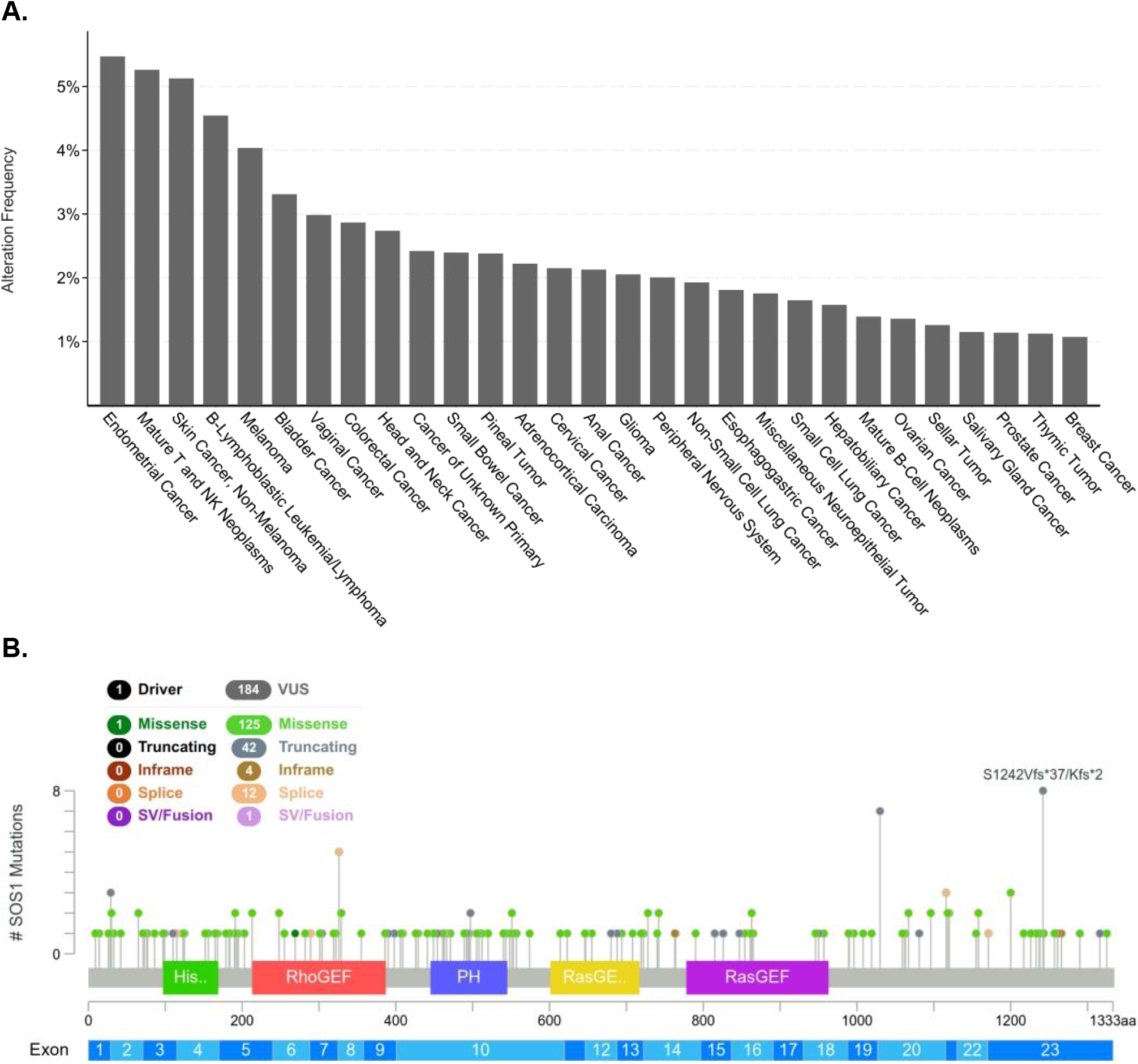
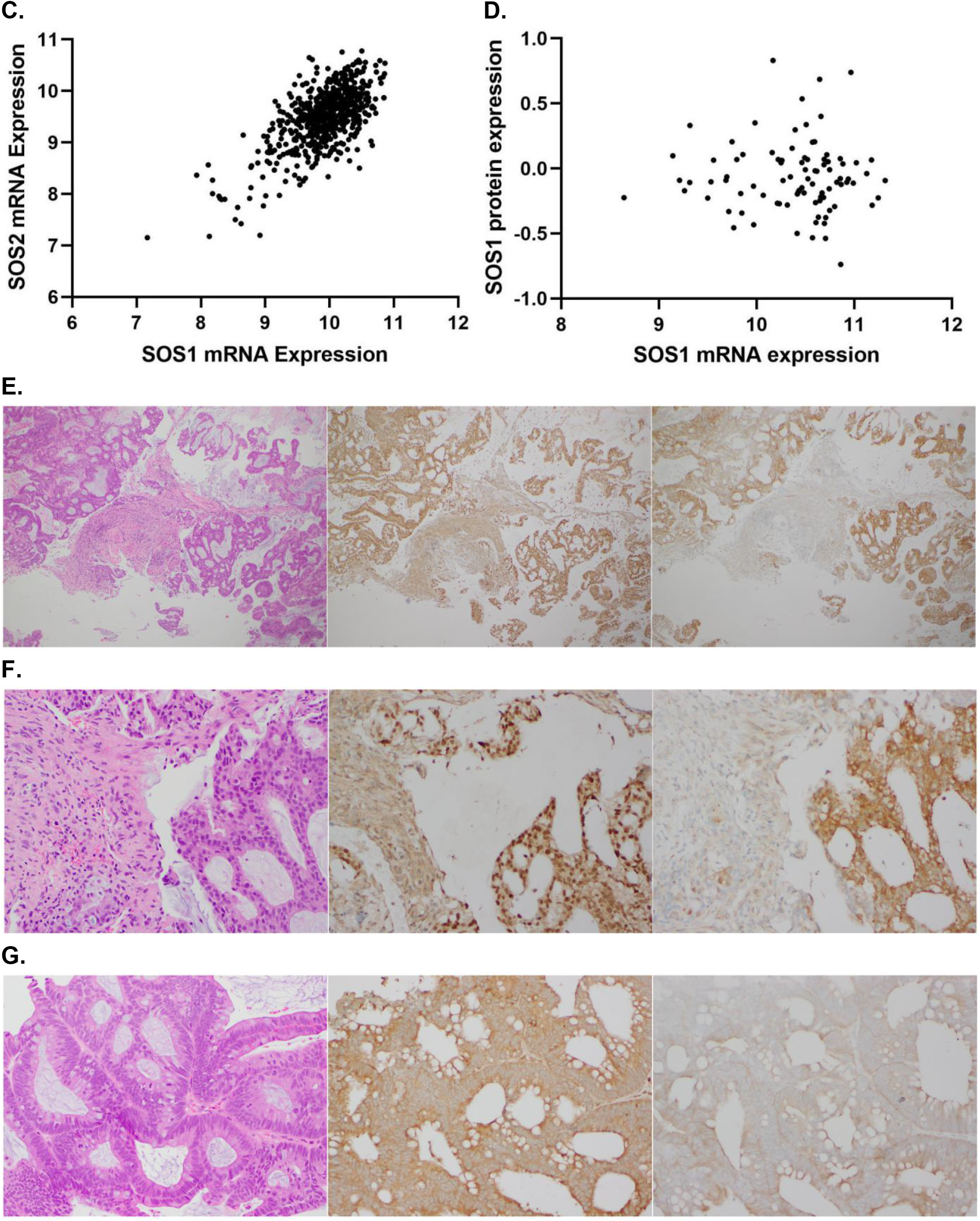
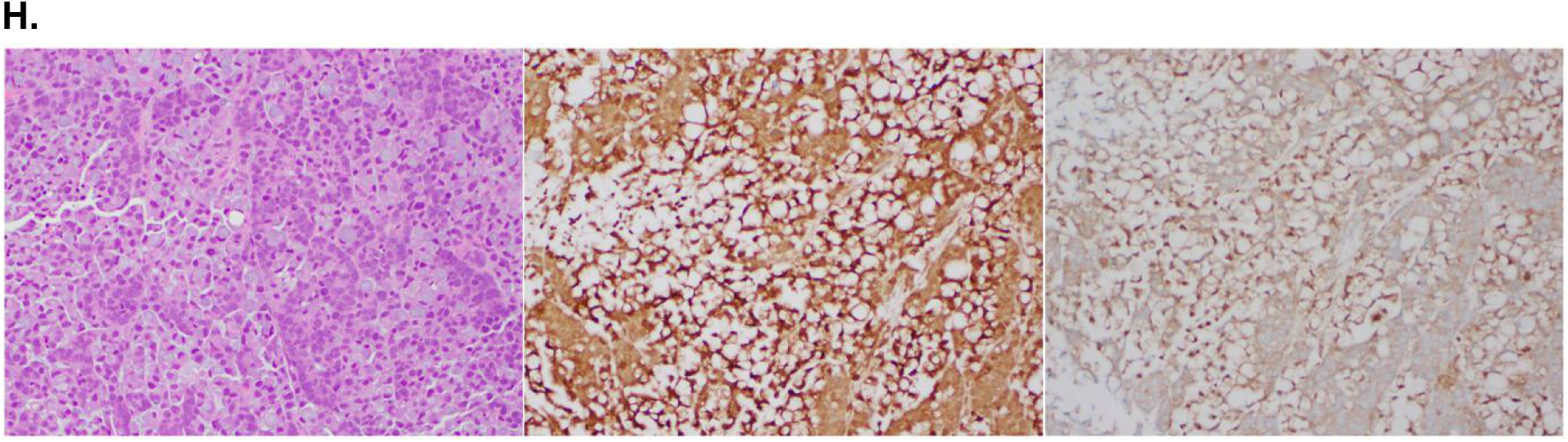
Molecular alterations, mRNA, and protein expressions of SOS1 in CRC. **(A)** Prevalence of *SOS1* mutations across different cancer types in GENIE cohort v11.0. **(B)** Distribution of *SOS1* alterations in CRC. NM_005633 | ENST00000402219 CCDS1802 | SOS1_HUMAN in GENIE cohort v11.0. Putative drivers versus variants of unknown significance was determined by OncoKB and hotspots. **(C)** Correlation between *SOS1* and *SOS2* mRNA expressions (RSEM, Batch normalized from Illumina HiSeq_RNASeqV2, log_2_) in CRC in TCGA PanCancer Atlas. mRNA expression as log_2_(value + 1), Spearman’s ρ: 0.56, *p*<0.001. **(D)** Correlation between *SOS1* mRNA (RSEM, Batch normalized from Illumina HiSeq_RNASeqV2, log_2_) and SOS1 protein expressions (mass spectrometry by CPTAC) in CRC in CPTAC-2 Prospective (Cell 2019). Spearman’s ρ: *-*0.05, *p*=0.6. **(E)** SOS1 and SOS2 expressions by IHC in CRC tissue in the low-power field. Left: H&E; Middle: SOS1 H-score 6; Right: SOS2 H-score 6. **(F)** SOS1 and SOS2 expressions by IHC in CRC tissue in the high-power field. Left: H&E; Middle: SOS1 H-score 6; Right: SOS2 H-score 6. **(G)** SOS1 and SOS2 expressions in moderately differentiated colon adenocarcinoma. Left: H&E; Middle: SOS1 H-score 6; Right: SOS2 H-score 3. **(H)** SOS1 and SOS2 expressions in poorly differentiated mucinous colon adenocarcinoma. Left: H&E; Middle: SOS1 H-score 9; Right: SOS2 H-score 6.

As to *SOS1* mRNA expression, there was no statistical difference between *RAS/RAF* wild-type and *RAS/RAF* mutant CRC (*p*=0.3) and across different tumor stages (*p*=1) in the TCGA PanCancer Altas (n=578) **(Supplemental Figure 1F-G)**. However, there was a significant correlation between *SOS1* and *SOS2* mRNA expression levels with Spearman’s ρ of 0.56 (*p*<0.001) **(Figure 2C)**. Similarly, SOS1 protein was not differentially expressed between *RAS/RAF* wild-type and mutant CRC (*p*=0.3) and across different tumor stages (*p*=0.4) in the CPTAC-2 CRC cohort (n=89) **(Supplemental Figure 1H-I)**. There was only a trend of correlation between SOS1 and SOS2 protein express levels (Spearman’s ρ 0.45, p=0.1) **(Supplemental Figure 1J)**, maybe partially due to small sample size or the lack of correlation between *SOS1* mRNA and SOS1 protein expression levels (Spearman’s ρ -0.05, p=0.6) **(Figure 2D)**. We further evaluated SOS1 and SOS2 protein expression by IHC in surgically resected CRC tissues. SOS1 and SOS2 were universally expressed in cancer cells with only minimal to no SOS1 and SOS2 expression in surrounding non-malignant tissues **(Figure 2E-F)**. SOS1 and SOS2 expressions patterns including nuclear, cytoplasmic and likely inner membranous expressions were also very different in morphologically distinct CRC specimens **(Figure 2G-H)**. In addition, the expression levels of SOS1 and SOS2 are not always correlated as assessed by IHC.

Further evaluation of SOS1 and SOS2 protein expression by IHC in CRC PDX models showed that their expression levels and SOS1/SOS2 protein expression ratio remained rather stable across PDX generational passages 1 through 3 **(Supplemental Figure 2A-C)**, between primary and metastatic CRC models **(Supplemental Figure 2D-F)**, and between *KRAS* wild-type and mutant CRC PDX models **(Supplemental Figure 2G-I)**.

### Prognostic value of SOS1 protein expression in CRC

We quantified our CRC PDX tumor growth using adjusted AUC (aAUC) and found that neither SOS1 or SOS2 protein expression levels nor SOS1/SOS2 protein expression ratio group (elevated if ratio>1, not elevated if ratio≤1) were associated with CRC PDX tumor growth **(Supplemental Figure 2J-L)**. Of note, CRC PDX tumor growth was not associated with *KRAS* mutation status (*p*=0.5) or primary versus metastatic CRC (*p*=0.8), either.

In contrast, in the CPTAC-2 CRC patient cohort, when SOS1 protein expression level measured by mass spectrometry was dichotomized using Z-score 0 as a cutoff, SOS1 high group (38 cases and 5 events) had significantly worse OS (*p*=0.048) compared to SOS1 low group (52 cases and 2 events) **(Figure 3A)**. Similarly, in patients with *RAS/RAF* mutant CRC, SOS1 high group has significantly worse OS (*p*=0.04) compared to SOS1 low group **(Figure 3B)**. However, when similar analyses were performed for *SOS1* mRNA expression, higher *SOS1* mRNA expression was not associated with OS in the entire cohort (*p*=0.2) or patients with *RAS/RAF* mutant CRC (*p*=0.051) **(Figure 3C-D)**.

**Figure 3.**
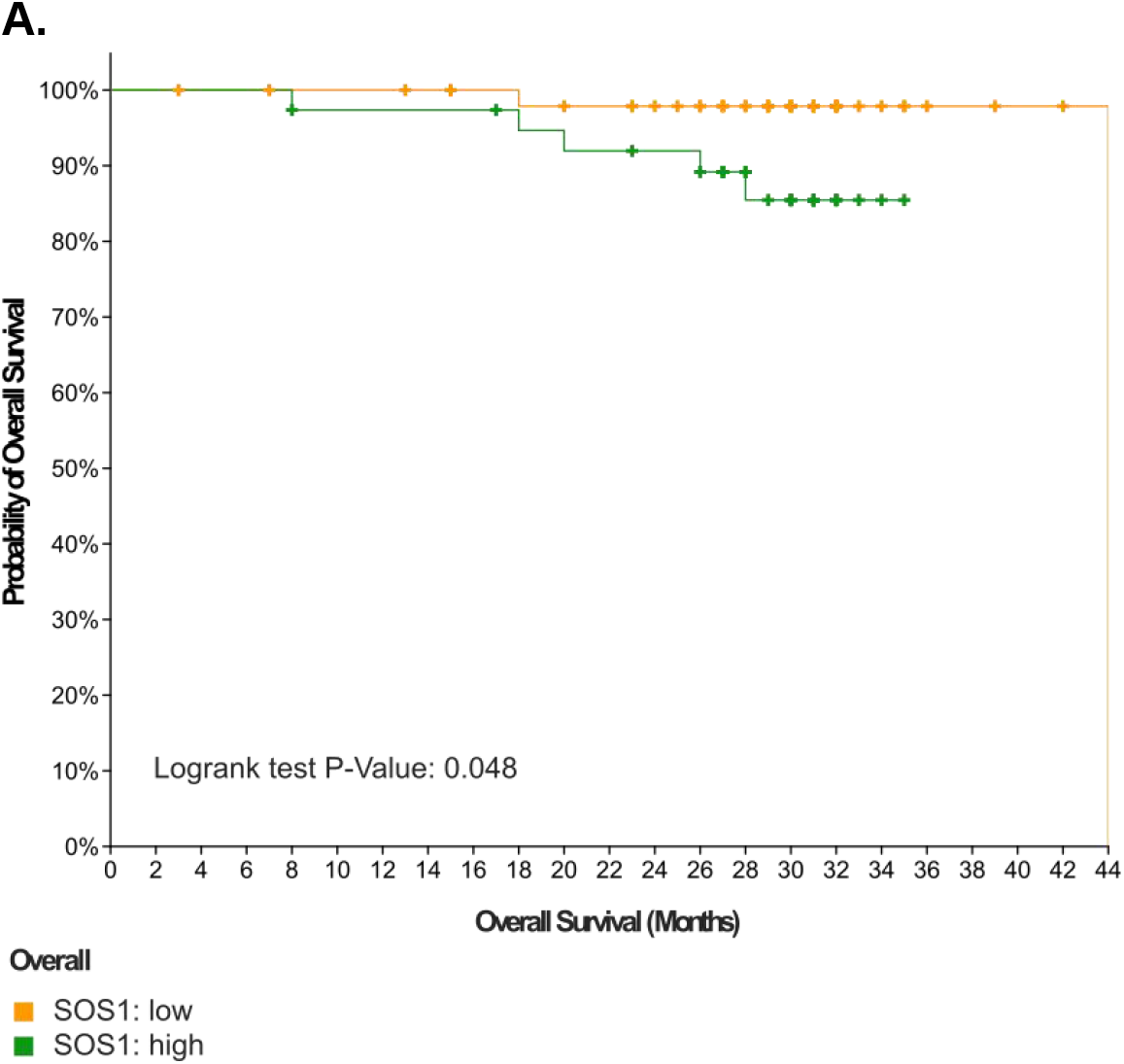
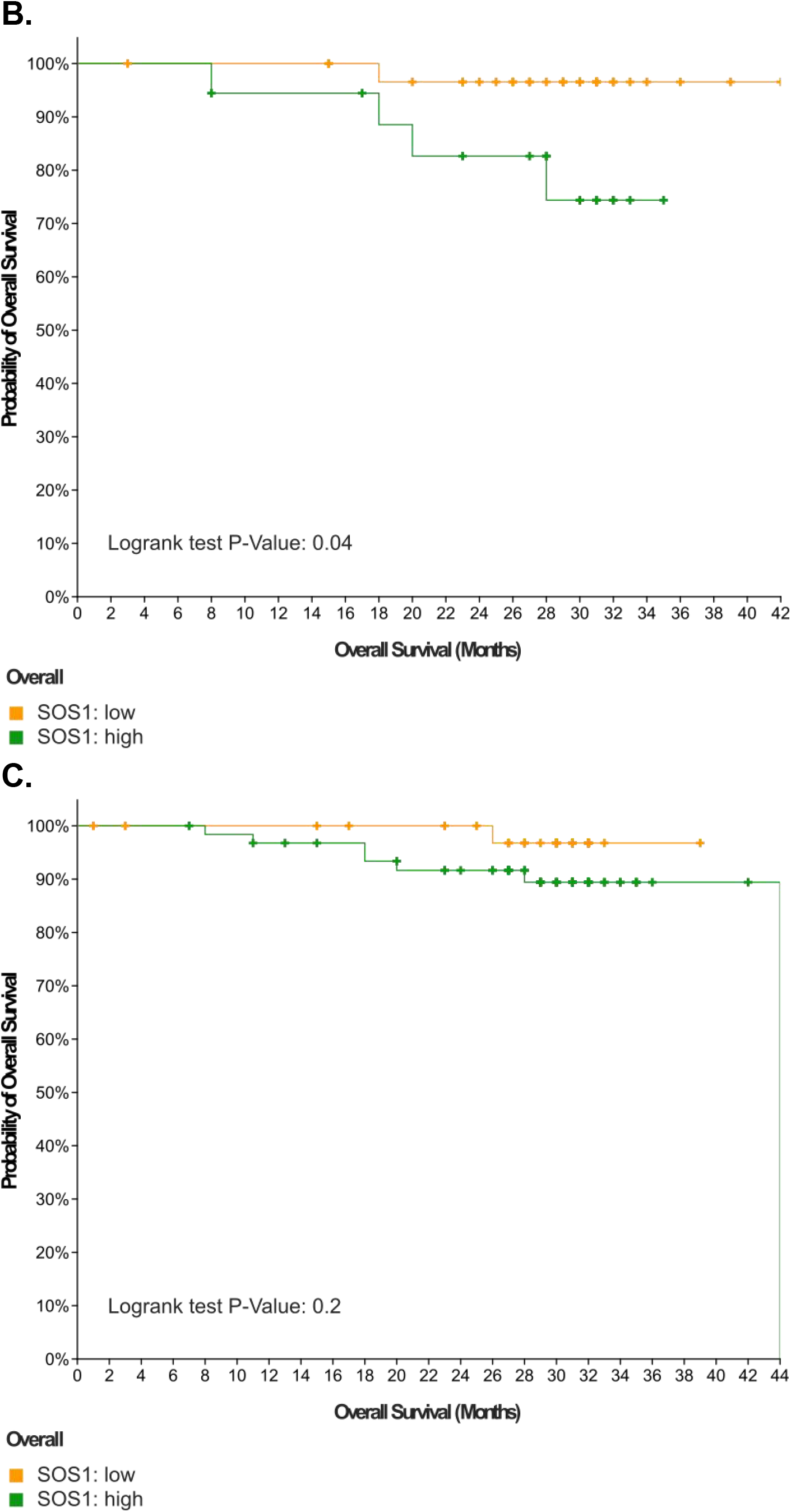
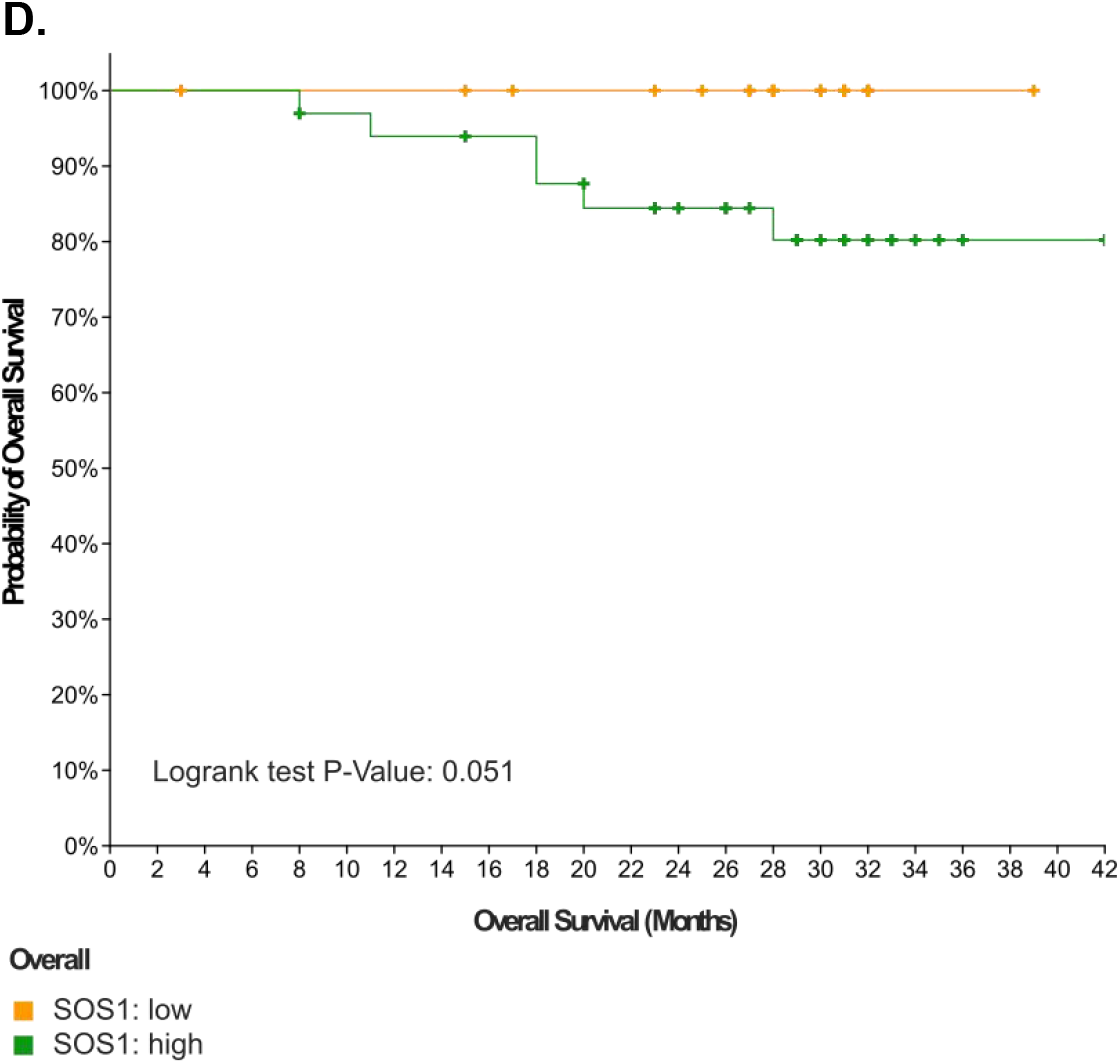
The prognostic value of *SOS1* mRNA and protein expressions in patients with CRC. OS in patients with CRC stratified by SOS1 protein expression (mass spectrometry by CPTAC) in CPTAC-2 Prospective. Z-score 0 was used as a cutoff. SOS1 low group had 52 cases with 2 events, median OS 44 months. SOS1 high group had 38 cases with 5 events. **(B)** OS in patients with *RAS/RAF* mutant CRC stratified by SOS1 protein expression (mass spectrometry by CPTAC) in CPTAC-2 Prospective. Z-score 0 was used as a cutoff. SOS1 low group had 32 cases with 1 event. SOS1 high group had 18 cases with 4 events. **(C)** OS in patients with CRC stratified by *SOS1* mRNA expression in CPTAC-2 Prospective. Z-scores relative to all samples (RNA Seq V2 RSEM UQ Log2) 0 was used as a cutoff. SOS1 low group had 37 cases with 1 event. SOS1 high group had 63 cases with 7 events, median OS 44 months. **(D)** OS in patients with *RAS/RAF* mutant CRC stratified by *SOS1* mRNA expression in CPTAC-2 Prospective. Z-scores relative to all samples (RNA Seq V2 RSEM UQ Log2) 0 was used as a cutoff. SOS1 low group had 21 cases with 0 events. SOS1 high group had 33 cases with 6 events.

### Predictive markers to SOS1 inhibitor sensitivity

We investigated potential predictive markers to SOS1 dependency as well as sensitivity to SOS1 inhibitor BI3406. The latter could be more clinically meaningful. Given previous report on the association between *KRAS G12/G13* mutations and BI3406 sensitivity, we treated our CRC PDO models with BI3406 as shown previously **(Figure 1C)** and found that CRC PDOs, either *KRAS* mutant or *KRAS* wild-type had responses to BI3406 equally (*p*=1) **(Figure 4A)**. In addition, interrogation of the DepMap database for the dependency of CRC cell lines to common genes in *KRAS* pathway showed that cells that are more dependent on *SOS1* do not have *KRAS* driver mutations **(Figure 4B)**. There was no correlation (Spearman’s ρ 0.18, *p*=0.2) between *SOS1* and *KRAS* dependency scores in 54 CRC cell lines **(Figure 4C)**. Further analysis of gene expressions of CRC cell lines with differential sensitivities (3 sensitive and 5 resistant) to BI3406 as defined by Hoffmann et al.^13^ identified gene features that were able to distinguish sensitive CRC cell lines from resistant ones **(Supplemental Figure 3A)**. GSEA identified 9 gene sets that are significantly enriched in BI3406-sensitive cell lines at nominal *p*<0.01 including Hallmark_E2F_Targets and Hallmark_MYC_Targets_V1 **(Supplemental Figure 3B)**. Similar observations were found in CRC cell lines with differential *SOS1* dependencies (8 dependent and 40 independent) as defined by CRISPR knockout experiments **(Supplemental Figure 3C)** along with 17 gene sets significantly enriched in *SOS1*-dependent CRC cell lines at nominal *p*<0.01 including Hallmark_Interferon_Alpha_Response and Hallmark_Interferon_Gamma_Response **(Supplemental Figure 3D)**. With further GSEA of CRC cell lines based on their *KRAS* dependency, we found that only a proportion of the enriched gene sets are shared among BI3406-sensitive, *SOS1* dependent, and *KRAS* dependent CRC cells **(Supplemental Figure 3E)**. In summary, *KRAS* mutations may not be a good predictive marker to either SOS1 inhibitor sensitivity or *SOS1* dependency (genetic knockout) in CRC.

**Figure 4.**
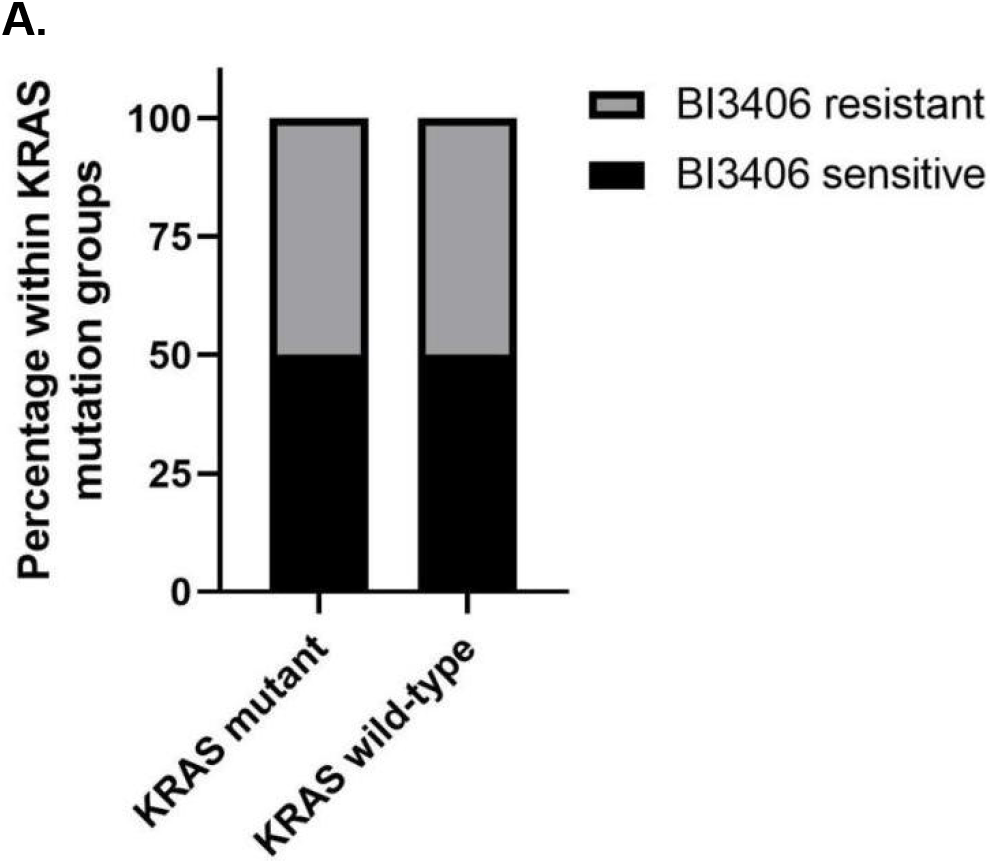
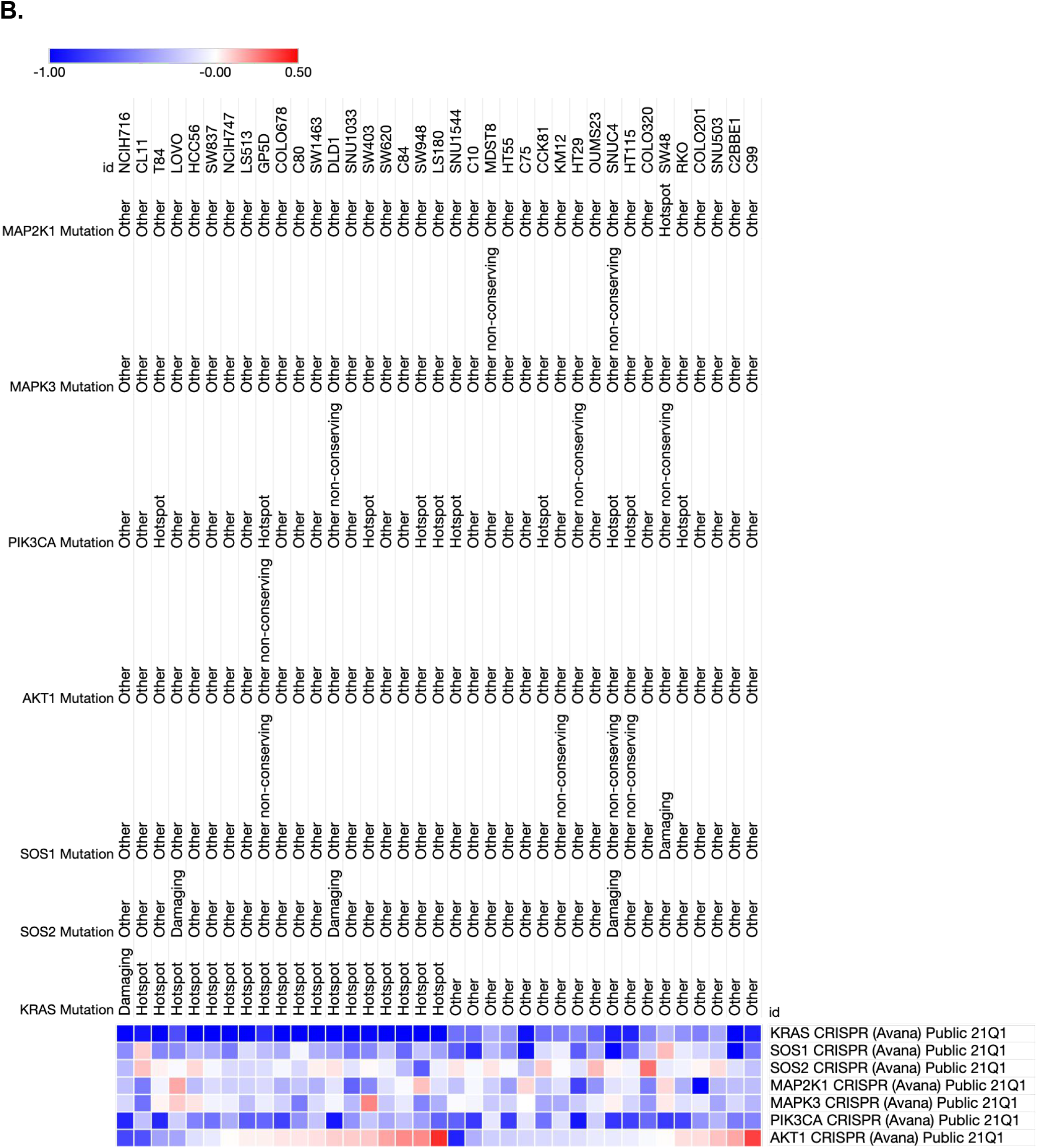
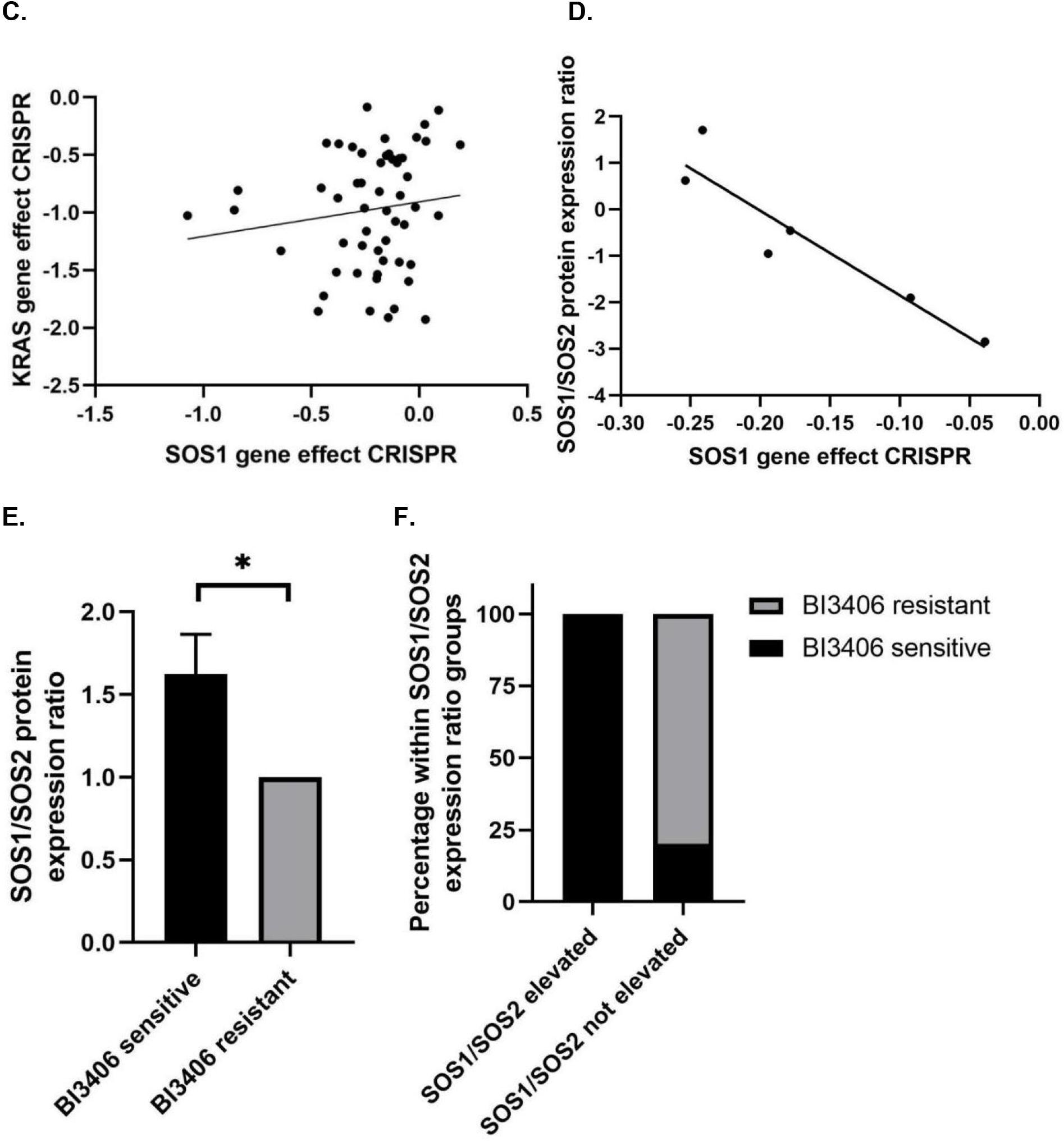
Predictive markers to *SOS1* dependency and sensitivity to SOS1 inhibitor BI3406 in CRC models. **(A)** Association of BI3406 sensitivity with *KRAS* mutation status in CRC PDO. *p*=1. Dependency of CRC cell lines to *SOS1* or *SOS2* knockout by CRISPR compared to other genes of *KRAS* signaling pathway in DepMap database. Dependency score less than zero suggests dependency of gene knockout. **(C)** Association of *SOS1* dependency with *KRAS* dependency in 54 CRC cell lines in DepMap database. Spearman’s ρ: 0.18, *p*=0.2. **(D)** Association of SOS1/SOS2 protein expression ratio with *SOS1* dependency in DepMap database. Spearman’s ρ: -0.886, *p*=0.007. **(E)** SOS1/SOS2 expression ratio by IHC in BI3406 sensitive and BI3406 resistant CRC PDXs. *p*=0.04. **(F)** The proportion of BI3406 sensitivity in SOS1/SOS2 expression ratio elevated and not elevated groups. The SOS1/SOS2 elevated group is defined as SOS1 H-score/SOS2 H-score >1. *p*=0.03.

We then evaluated the potential of SOS1 and SOS2 protein expressions as predictive markers. We found that higher SOS1/SOS2 protein expression ratio in CRC cell lines measured by mass spectrometry was significantly associated *SOS1* dependency (Spearman’s ρ -0.886, *p*=0.007) **(Figure 4D)** instead of SOS1 protein expression, *SOS1* or *SOS2* mRNA expression or their ratio **(Supplemental Figure 4A-C)**. Higher SOS1/SOS2 protein expression ratio by IHC in our CRC PDOs predicted sensitivity to BI3406 (*p*=0.04) **(Figure 4E)**. Practically, all cases in SOS1/SOS2 expression ratio elevated group defined as SOS1/SOS2 H-score ratio>1 were sensitive to SOS1 inhibition by BI3406 (*p*=0.03) **(Figure 4F)**. Neither SOS1 or SOS2 protein expression alone was associated with BI3406 sensitivity **(Supplemental Figure 4D-E)**.

We then compared gene expressions between BI3406-sensitive CRC PDOs (MCC19990-006, MCC19990-010, and MCC19990-013) and resistant CRC PDOs (MCC19990-002, MCC19990-007, and MCC20584-010). We identified a number of genes that were differentially expressed in these CRC PDOs which may be further evaluated for their predictive value to SOS1 sensitivity **(Figure 5A)**. GSEA identified 7 gene sets significant enriched at nominal *p*<0.01 in BI3406 resistant CRC PDOs **(Figure 5B)** with the top three including Hallmark_Cholesterol_Homeostasis, Hallmark_Epithelial_Mesenchymal_Transition, and Hallmark_TNFA_Signaling_Via_NFκB **(Figure 5C)**.

**Figure 5.**
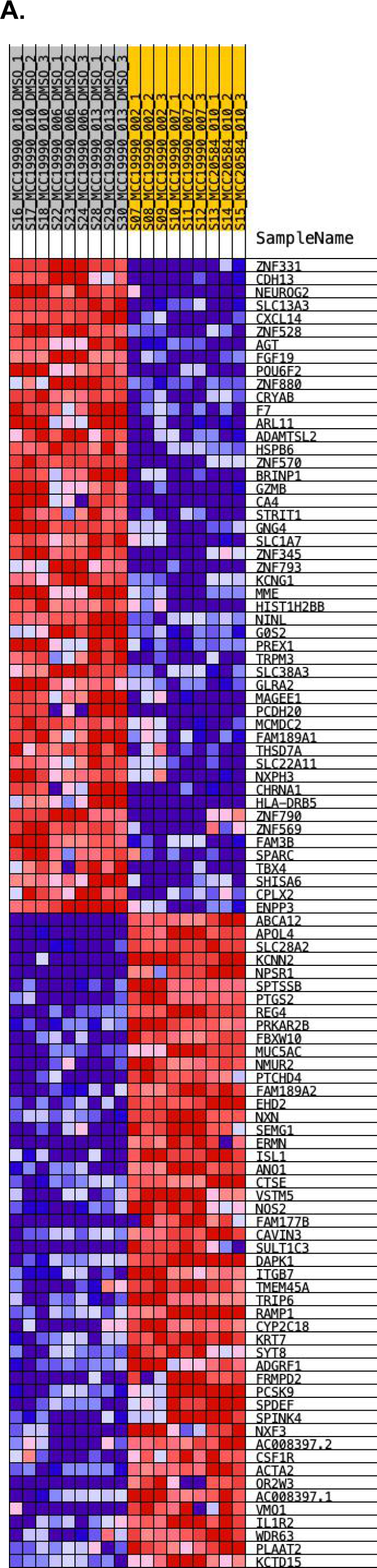
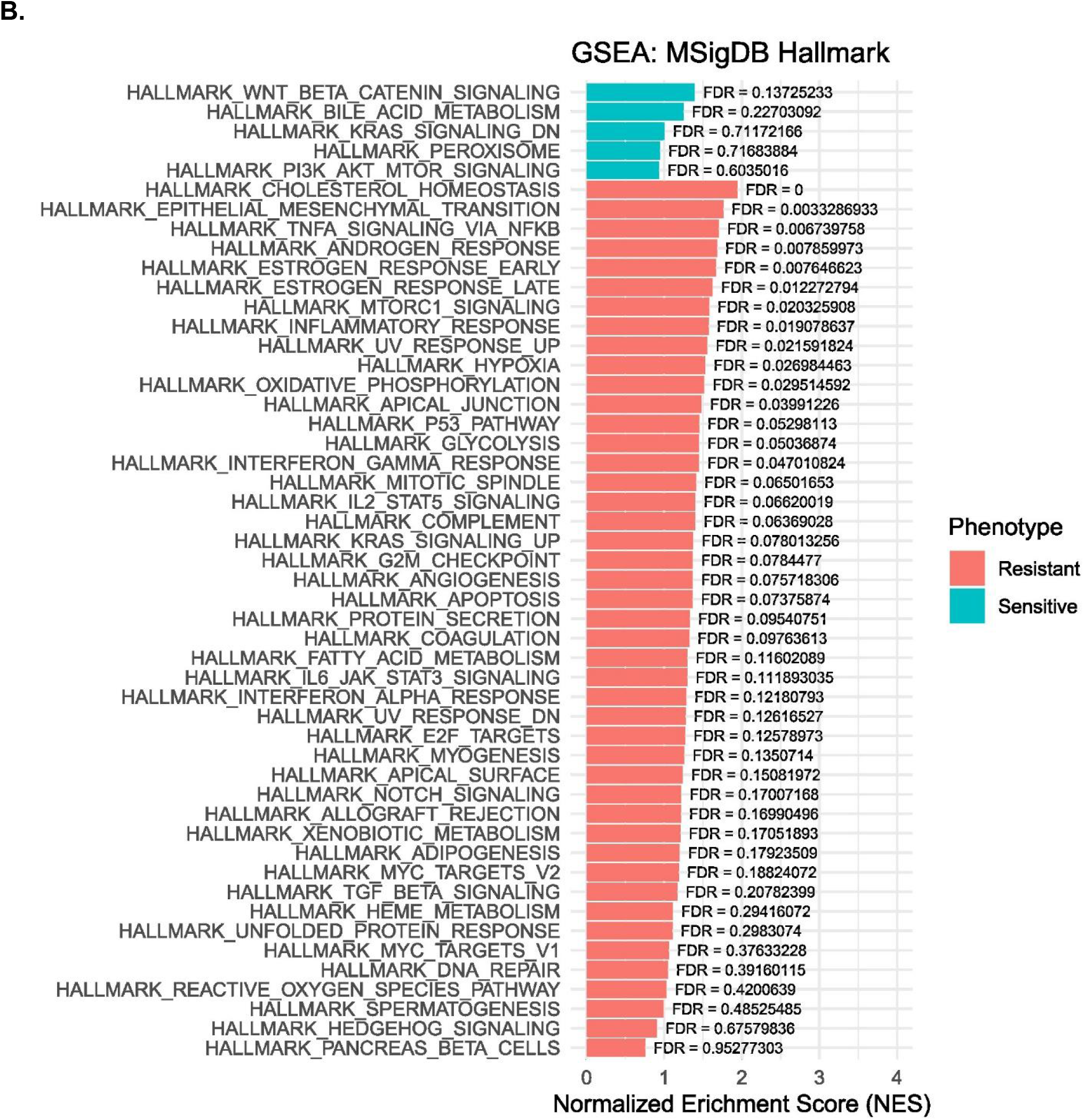
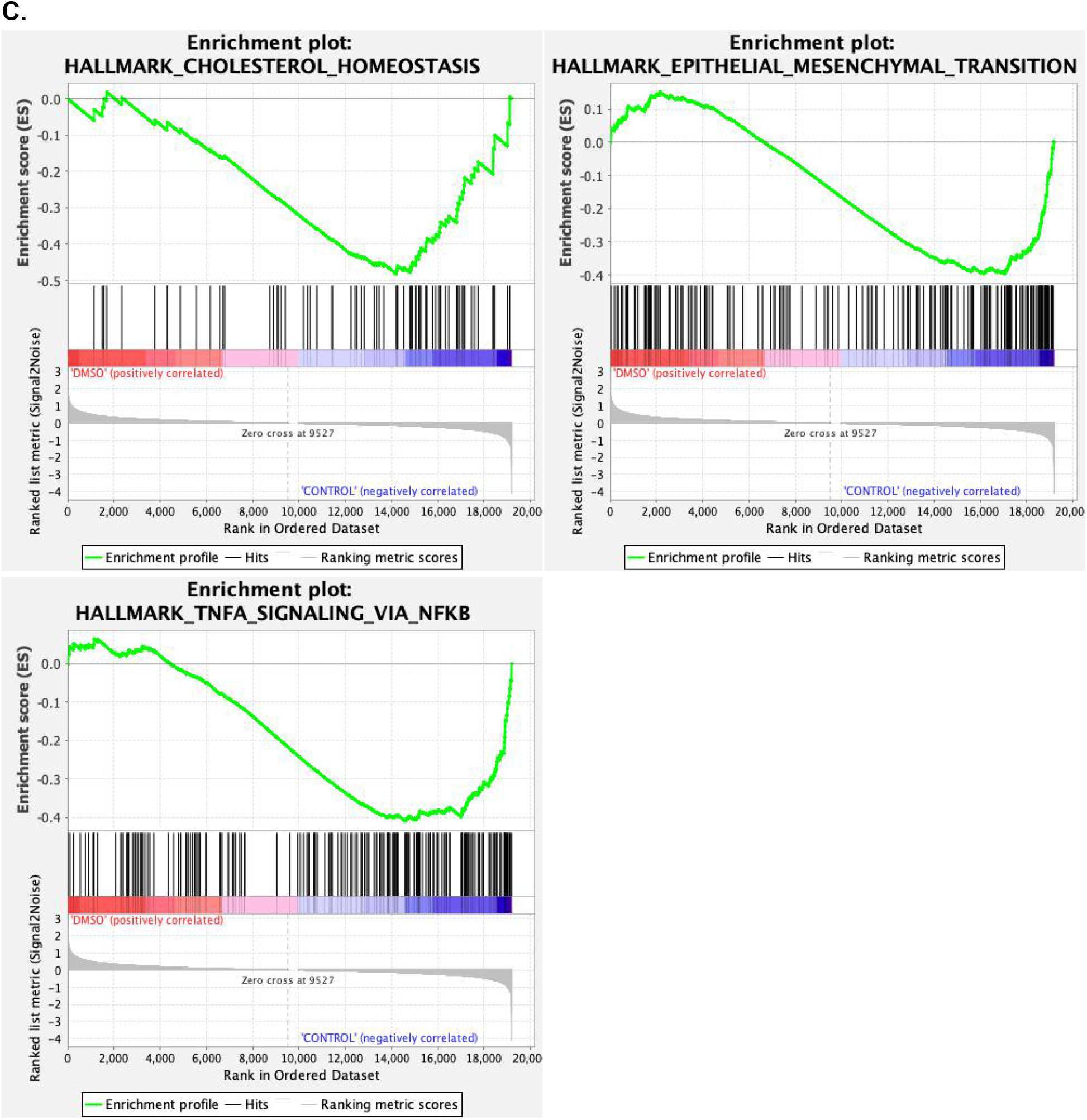
mRNA expression and gene set enrichment analyses of CRC PDOs. **(A)** Gene expressions of CRC PDOs with differential sensitivity to SOS1 inhibitor BI3406. The dataset has 19195 features (genes). # of markers for phenotype sensitive: 9527 (49.6%) with correlation area 50.3%. # of markers for phenotype resistant: 9668 (50.4%) with correlation area 49.7%. Heat Map included the top 50 features for each phenotype sensitive (n=3) or resistant (n=3) to BI3406. **(B)** Summary of GSEA of CRC PDOs with differential sensitivity to SOS1 inhibitor BI3406. **(C)** Enrichment plots in BI3406 sensitive vs resistant CRC PDOs. 39/50 gene sets are upregulated in phenotype resistant. 13 gene sets are significant enriched at FDR <25%. 7 gene sets are significantly enriched at nominal p value <1%. 11 gene sets are significantly enriched at nominal p value < 5%. 11/50 gene sets are upregulated in phenotype sensitive. 0 gene set is significantly enriched at FDR < 25%. 0 gene set is significantly enriched at nominal p value < 1%. 1 gene set is significantly enriched at nominal p value < 5%. Snapshots of enrichment results in resistant CRC PDOs are shown.

### Cellular effect of SOS1 inhibition in CRC

To further understand the findings on predictive markers to SOS1 inhibitor sensitivity, we evaluated the cellular effect of SOS1 inhibition in CRC. We found that phosphorylation of ERK (pERK) appeared to be moderately suppressed by BI3406 in a concentration-dependent manner in HCT116 **(Supplemental Figure 5A)**. However, the GTP-bound RAS level had a clear rebound at 48 hours despite various levels of initial suppression upon treatment with BI3406 in both SOS1-inhibitor resistant (MCC19990-002, MCC19990-007) and sensitive (MCC19990-010) CRC PDOs **(Figure 6A)**. Expressions of the 9 *KRAS* effector genes in the 3 BI3406-sensitive CRC PDOs after treatment with BI3406 for 24 hours did not show significant difference **(Figure 6B)** despite suppression of some enriched gene sets such as Hallmark_MYC_Targets_V1 and Hallmark_E2F_Targets **(Supplemental Figure 5B)**, concordant with the observations in sensitive CRC cell lines **(Supplemental Figure 3B)**.

**Figure 6.**
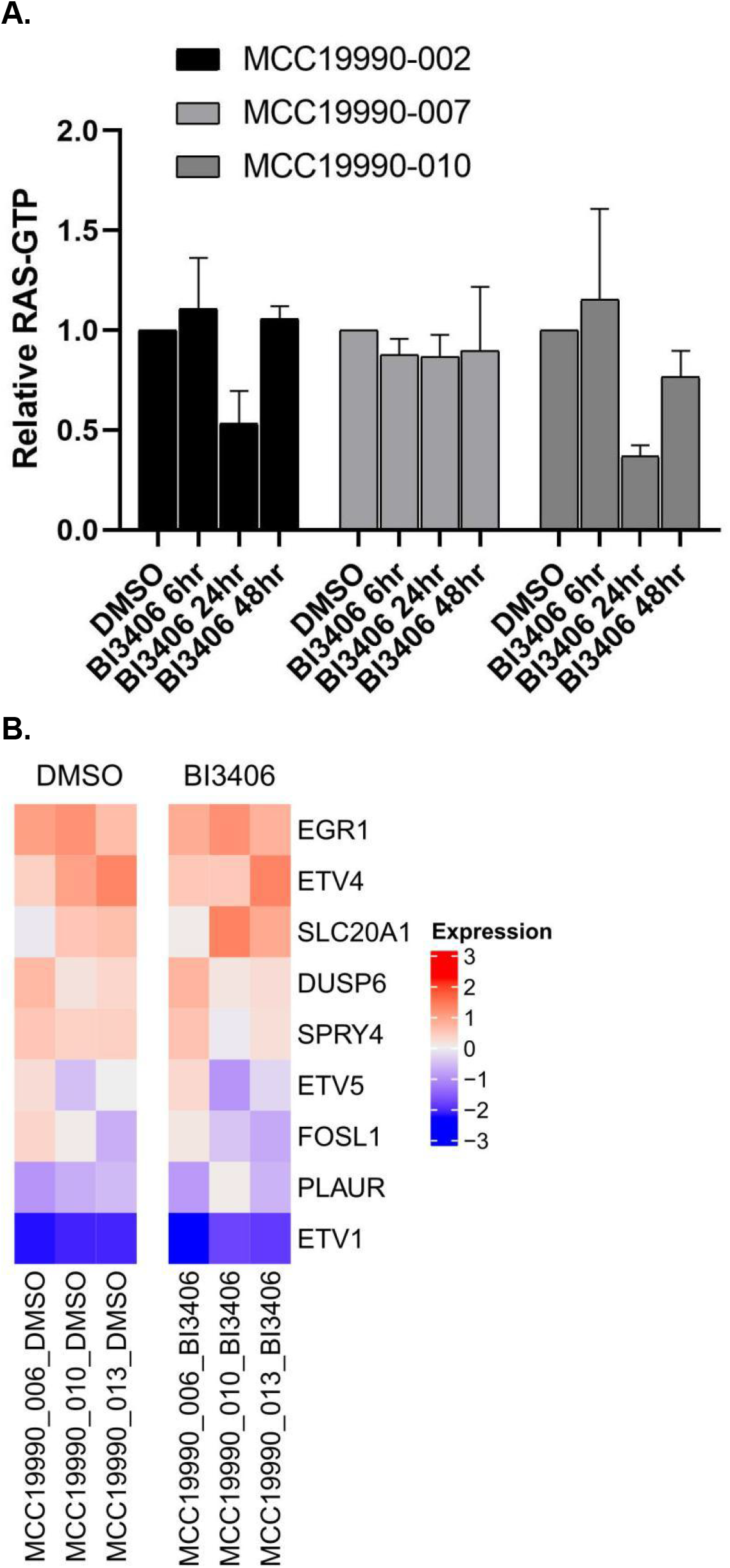
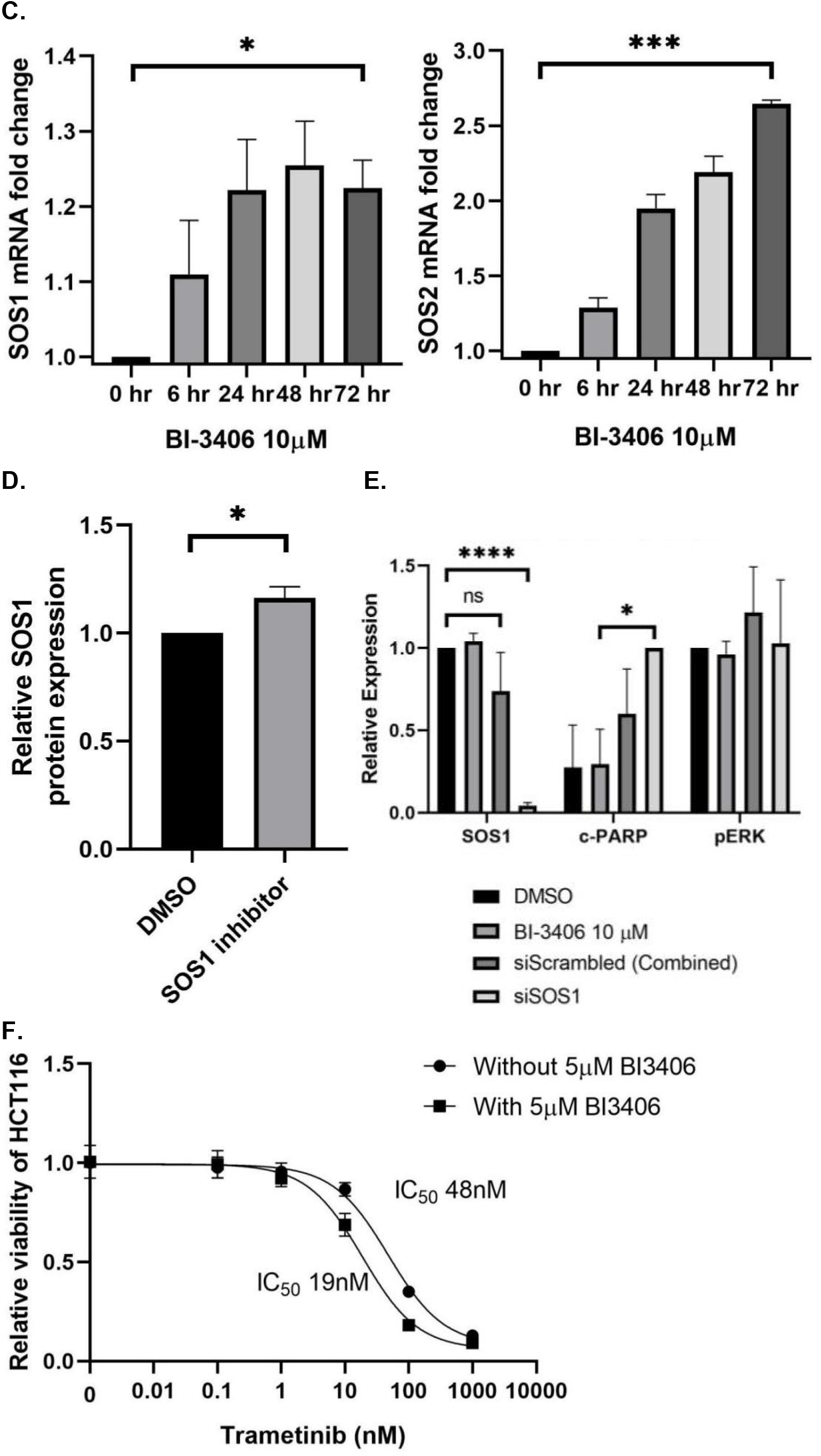
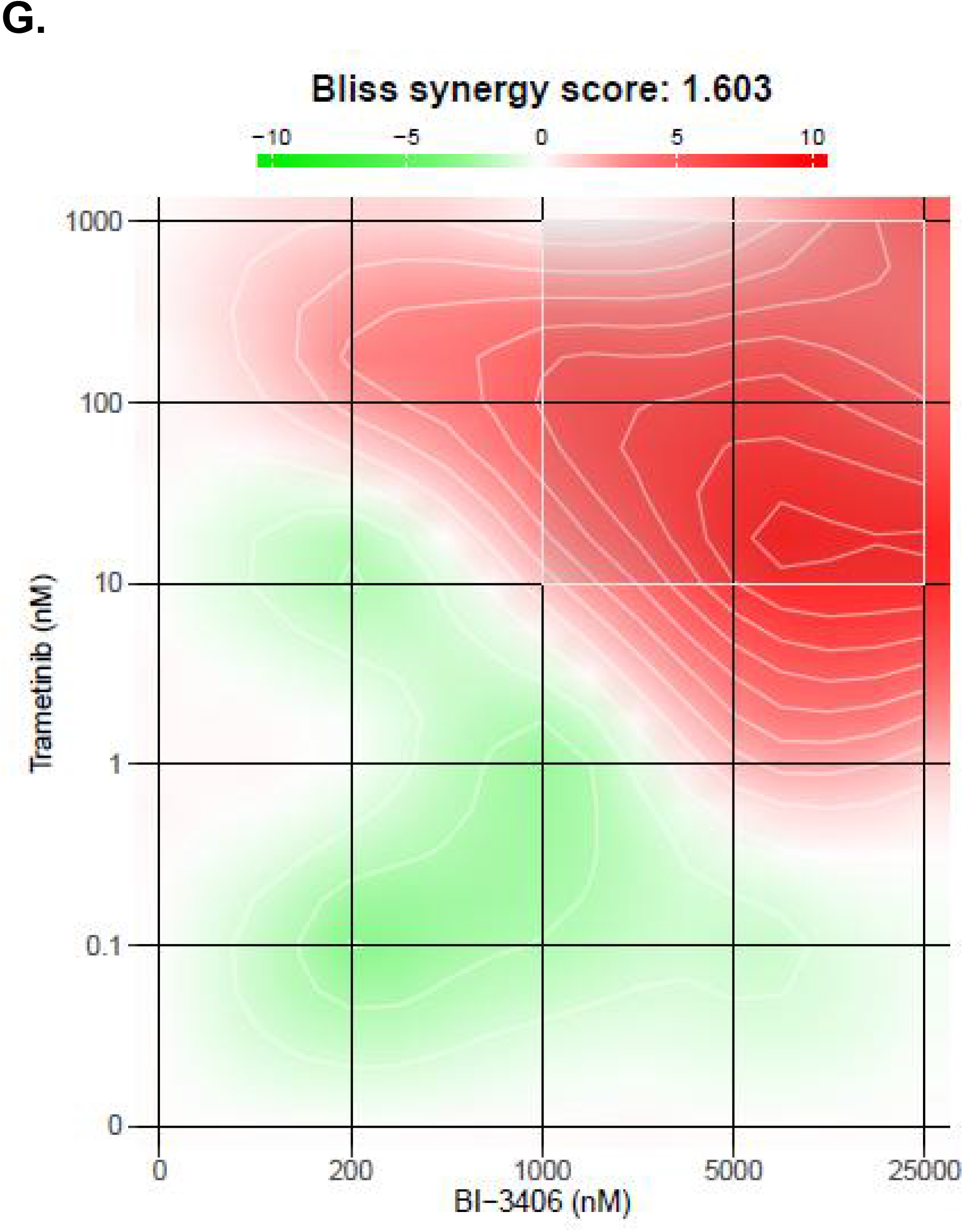
Effect of SOS1 inhibitor BI3406 on CRC cells and PDOs. **(A)** GTP-bound KRAS levels in CRC PDOs after treatment with SOS1 inhibitor BI3406 at 0, 6, 24, and 48 hours. **(B)** Expression of 9 *KRAS* effector genes in BI3406 sensitive CRC PDOs before and after treatment with 1µM BI3406 for 24 hours. **(C)** *SOS1* and *SOS2* mRNA expression levels in HCT116 after treatment with 10µM BI3406 for 0, 6, 24, 48 and 72 hours. *p*=0.03 for *SOS1* mRNA and *p*<0.0001 for *SOS2* mRNA. **(D)** SOS1 protein expression in HCT116 after treatment of 10µM SOS1 inhibitor (BI68BS or BI3406) for 6 hours. *p*=0.01. **(E)** *SOS1* knockdown with siRNA after 48 hours was associated with increased apoptosis in SOS1 inhibitor BI3406 resistant CRC cell line HCT116. **(F)** Relative viability of HCT116 after treatment with various concentrations of trametinib in the presence or absence of BI3406 for 72 hours. **(G)** BI3406 and trametinib have synergistic cytotoxic effects in HCT116 as estimated using Bliss model.

To reveal potential cellular adaptation mechanisms, we measured *SOS1* and *SOS2* mRNA expression levels and found that both levels significantly increased upon treatment with BI3406 for up to 72 hours (*p*=0.03 for *SOS1* mRNA, *p*<0.0001 for *SOS2* mRNA) **(Figure 6C)**. The same adaption was found for the significant increase of SOS1 protein expression (*p*=0.01) after treatment with SOS1 inhibitors for only 6 hours **(Figure 6D)**.

To assess potential strategies that may overcome these mechanisms, we effectively knocked down SOS1 with siRNA. When compared with SOS1 inhibition by BI3406, we observed significantly increased cleaved PARP despite the lack of pERK suppression as shown in **Figure 6E** and **Supplemental Figure 5C**. We also found that the presence of BI3406 can significantly decrease the IC_50_ of MEK inhibitor trametinib from 48nM to 19nM in HCT116 (*p*<0.001) **(Figure 6F)**. BI3406 and trametinib had synergistic cytotoxic effects in HCT116 with Bliss synergy score of 1.6 **(Figure 6G)**. In contrast, BI3406 was not synergistic with pan-class I PI3K inhibitor buparlisib or FAK inhibitor defactinib **(Supplemental Figure 5D-E)**.

## Discussion

In this study, we aimed to evaluate the translational role of SOS1 in CRC. We first made an effort to establish dedicated preclinical CRC models, which can demonstrate differential sensitivities to available SOS1 inhibitor BI3406. However, although we obtained dose-response curves in various CRC cell lines similar to those previously reported in the discovery of BI3406,^13^ yet the range of IC_50_ was narrow with the lowest IC_50_ of one of our primary CRC cell lines close to 10µM. This is concerning for drug discovery efforts moving forward due to possible off-target effects from high drug concentration. Increased number of various somatic mutations in commercial cell lines may increase the chance of resistance to small molecules and provide false negative findings compared to patient-derived cancer models where the pattern of somatic mutations and other molecular alterations are more clearly defined.^20^ Consistent with previous findings on CRC PDOs, which can recapitulated disease biology and patient’s response to therapy,^23,32^ our CRC PDOs had much better differential sensitivities to BI3406 with wider IC_50_ range compared to CRC cell lines. These CRC PDOs served as excellent models for subsequent translational research studies on SOS1.

*SOS1* activating mutations have been reported in Noonan’s syndrome and other RASopathies.^47–49^ However, driver mutations were reported to be rare similar to what we found in CRC.^50^ Therefore, *SOS1* mutations are unlikely to be responsible for the pathogenesis of a subset of CRC or used as prognostic or predictive markers. The findings on the positive correlation between SOS1 and SOS2 mRNA and protein expressions supported previous hypotheses on similar GEF function for RTK-dependent RAS activation by SOS2. SOS2 may be more related to PI3K signaling instead of MAPK signaling. The latter is more regulated by SOS1.^19,51^ However, no SOS2 inhibitor is available to date despite that SOS2 inhibition may be more effective in suppression of *RTK/RAS* signaling.^19,51^ In an effort to tease out the relationship between SOS1 and SOS2 protein expressions, we immunochemically stained them in specimens from patients with CRC. Universal expression in tumor tissue along with minimal expression in surround normal tissue supported that SOS1 has the potential to be a selective target in CRC sparing the on and off target toxicities in normal tissue. However, we also noticed that the patterns and levels of SOS1 and SOS2 expressions in CRC with different morphologies are highly heterogeneous which warranted further investigations on their functional significance in future studies with larger sample size. SOS1 and SOS2 protein expressions by IHC were fairly stable without significant changes across CRC PDX passages, histopathologic, and genetic variables, which supported their potential as robust biomarkers. Higher SOS1 protein expression as a poor prognostic marker for OS in patients with CRC provided the rationale to target SOS1 for inhibition or degradation.

The development of companion predictive biomarker has played critical role in the clinical success of targeted therapy and optimal patient selection in key clinical trials. In our effort to discover predictive biomarkers to the sensitivity of SOS1 inhibitor, we surprisingly did not find *KRAS G12/13* mutations as a predictive marker to SOS1 inhibition or dependency in CRC. This could be due to a number of reasons such as different tumor types studied, different tumor models used, the presence of other co-mutations as reported previously,^13,52^ and off-target activities of SOS1 inhibitor used etc. The latter was supported by the fact that different gene sets were enriched among BI3406-sensitive, *SOS1*-dependent, and *KRAS*-dependent CRC cell lines. Gratifyingly, we found that SOS1/SOS2 protein expression ratio by IHC in CRC tissues was correlated with BI3406 sensitivity of CRC PDOs derived from the same tissue. This finding was independently confirmed by DepMap data showing SOS1/SOS2 protein expression ratio by mass spectrometry was highly correlated with *SOS1* dependency in CRC cell lines. This observation was in concordant with the role of SOS2 as a compensation mechanism to suppression of SOS1 function. In cases where SOS2 protein was less expressed or knocked down, the cancer cells became more sensitive to SOS1 inhibition.^13^ The comparison of mRNA expressions in SOS1 inhibition sensitive and resistant CRC PDOs revealed distinct genes and gene sets which could serve as additional predictive markers for BI3406 sensitivity or primary resistance mechanisms which could provide insights in combination therapy development. The top three gene sets enriched in BI3406-resistant CRC PDOs including cholesterol homeostasis,^53^ epithelial mesenchymal transition^54^ and TNFα/NFκB^55^ were all known to be related to the pathogenesis, progression, and resistance to therapy in CRC.

Our study on the cellular effect of BI3406 in CRC revealed potential cellular adaptation mechanisms. Rebound of GTP-bound RAS level at 48 hours upon treatment with BI3406 in not only resistant but also sensitive PDO models suggested that in resistant PDOs, inhibition of GEF function and downstream signaling could either be due to compensation from intrinsically activated alternative pathways such as those revealed above in the RNA-seq analysis, or from feedback upregulation of SOS1/2 expressions as observed in our experiments; In contrast, in sensitive PDOs, BI3406 induced antiproliferative effect at least may partially be due to off target activities such as those targets in the *MYC* and *E2F* pathways instead of *RTK/RAS* pathway. All of these findings could be hypothesis generating and thus warrant dedicated studies. To potentially overcome adaptive upregulation of SOS1 protein, genetic knockdown of SOS1 induced increased apoptosis of a BI3406-resistant CRC cell line, which opened the door for the use of pharmacological degradation of SOS1 to improve activity of targeting SOS1. On the other hand, rationale combination therapy strategies should be extensively explored with known synergy between BI3406 and MEK inhibitor but should be expanded to agents targeting pathways other than *RTK/RAS*.

In contrast to previous studies, our study utilized patient-relevant PDO models to study the role of SOS1, specifically in CRC. We identified novel and rationale predictive biomarkers to the sensitivity of SOS1 inhibition and provided information on potential cellular adaptation mechanisms. These findings although may be important for future clinical development of SOS1-targeting agents, yet to overcome the limitations of our study, rigorous validations of these findings are required in patient-derived *in vivo* models or in correlative studies of clinical trials with large sample size that is statistically powered for biomarker discovery.

In summary, our study suggested that CRC PDOs could serve as better models for translational study of SOS1 in CRC. High SOS1 protein expression was a worse prognostic marker in CRC. High SOS1/SOS2 protein expression ratio predicted sensitivity to SOS1 inhibition and dependency. Our preclinical findings supported further clinical development of SOS1-targeting agents in CRC.

## Author contributions

Hao Xie, Aik Choon Tan, and Jason B. Fleming designed this study. Diego Alem, Xinrui Yang, Francisca Beato, Bhaswati Sarcar, Alexandra F. Tassielli, Ruifan Dai, and Tara L. Hogenson performed the experiments. Kun Jiang, Francisca Beato, Ruifan Dai assisted in sample collection and pathology review. Hao Xie and Margaret A. Park analyzed the data. Hao Xie wrote the manuscript. Jianfeng Cai, Yu Yuan, Martin E. Fernandez-Zapico, Aik Choon Tan, Jason B. Fleming, and Hao Xie helped revision of the manuscript, funding acquisition, and project administration. All authors have read and approved the final manuscript.

## Acknowledgements

This study was supported by the Moffitt Cancer Center Support Grant to H. X., the Tissue Core, the Molecular Genomics Core, and the Biostatistics and Bioinformatics Shared Resource at the H. Lee Moffitt Cancer Center & Research Institute, an NCI designated Comprehensive Cancer Center (P30-CA076292).

## Data availability statement

The data of this study are available from the corresponding author upon reasonable request.

## References

1. Lakatos G, Köhne CH, Bodoky G. Current therapy of advanced colorectal cancer according to RAS/RAF mutational status. Cancer Metast Rev. Published online 2020:1–15. doi:10.1007/s10555-020-09913-7

2. Uprety D, Adjei AA. KRAS: From undruggable to a druggable Cancer Target. Cancer Treat Rev. 2020;89:102070. doi:10.1016/j.ctrv.2020.102070

3. Moore AR, Rosenberg SC, McCormick F, Malek S. RAS-targeted therapies: is the undruggable drugged? Nat Rev Drug Discov. 2020;19(8):533–552. doi:10.1038/s41573-020-0068-6

4. Freedman TS, Sondermann H, Friedland GD, et al. A Ras-induced conformational switch in the Ras activator Son of sevenless. Proc National Acad Sci. 2006;103(45):16692–16697. doi:10.1073/pnas.0608127103

5. Fedele C, Li S, Teng KW, et al. SHP2 inhibition diminishes KRASG12C cycling and promotes tumor microenvironment remodeling. J Exp Med. 2020;218(1):e20201414. doi:10.1084/jem.20201414

6. Abbott JR, Hodges TR, Daniels RN, et al. Discovery of Aminopiperidine Indoles That Activate the Guanine Nucleotide Exchange Factor SOS1 and Modulate RAS Signaling. J Med Chem. 2018;61(14):6002–6017. doi:10.1021/acs.jmedchem.8b00360

7. Abbott JR, Patel PA, Howes JE, et al. Discovery of Quinazolines That Activate SOS1-Mediated Nucleotide Exchange on RAS. Acs Med Chem Lett. 2018;9(9):941–946. doi:10.1021/acsmedchemlett.8b00296

8. Sarkar D, Olejniczak ET, Phan J, et al. Discovery of Sulfonamide-Derived Agonists of SOS1-Mediated Nucleotide Exchange on RAS Using Fragment-Based Methods. J Med Chem. 2020;63(15):8325–8337. doi:10.1021/acs.jmedchem.0c00511

9. Akan DT, Howes JE, Sai J, et al. Small Molecule SOS1 Agonists Modulate MAPK and PI3K Signaling via Independent Cellular Responses. Acs Chem Biol. 2019;14(3):325–331. doi:10.1021/acschembio.8b00869

10. Hodges TR, Abbott JR, Little AJ, et al. Discovery and Structure-Based Optimization of Benzimidazole-Derived Activators of SOS1-Mediated Nucleotide Exchange on RAS. J Med Chem. 2018;61(19):8875–8894. doi:10.1021/acs.jmedchem.8b01108

11. Burns MC, Sun Q, Daniels RN, et al. Approach for targeting Ras with small molecules that activate SOS-mediated nucleotide exchange. Proc National Acad Sci. 2014;111(9):3401–3406. doi:10.1073/pnas.1315798111

12. Hillig RC, Sautier B, Schroeder J, et al. Discovery of potent SOS1 inhibitors that block RAS activation via disruption of the RAS–SOS1 interaction. Proc National Acad Sci. 2019;116(7):2551–2560. doi:10.1073/pnas.1812963116

13. Hofmann MH, Gmachl M, Ramharter J, et al. BI-3406, a Potent and Selective SOS1–KRAS Interaction Inhibitor, Is Effective in KRAS-Driven Cancers through Combined MEK Inhibition. Cancer Discov. 2021;11(1):142–157. doi:10.1158/2159-8290.cd-20-0142

14. Ramharter J, Kessler D, Ettmayer P, et al. One Atom Makes All the Difference: Getting a Foot in the Door between SOS1 and KRAS. J Med Chem. Published online 2021. doi:10.1021/acs.jmedchem.0c01949

15. He H, Zhang Y, Xu J, et al. Discovery of Orally Bioavailable SOS1 Inhibitors for Suppressing KRAS-Driven Carcinoma. J Med Chem. Published online 2022. doi:10.1021/acs.jmedchem.2c00986

16. Ketcham JM, Haling J, Khare S, et al. Design and Discovery of MRTX0902, a Potent, Selective, Brain-Penetrant, and Orally Bioavailable Inhibitor of the SOS1:KRAS Protein–Protein Interaction. J Med Chem. Published online 2022. doi:10.1021/acs.jmedchem.2c00741

17. https://clinicaltrials.gov. A Study to Test Different Doses of BI 1701963 Alone and Combined With Trametinib in Patients With Different Types of Advanced Cancer (Solid Tumours With KRAS Mutation). Accessed July 6, 2022. https://clinicaltrials.gov/ct2/show/NCT04111458

18. Theard PL, Sheffels E, Sealover NE, Linke AJ, Pratico DJ, Kortum RL. Marked synergy by vertical inhibition of EGFR signaling in NSCLC spheroids shows SOS1 is a therapeutic target in EGFR-mutated cancer. Elife. 2020;9:e58204. doi:10.7554/elife.58204

19. Sheffels E, Kortum RL. Breaking Oncogene Addiction: Getting RTK/RAS-Mutated Cancers off the SOS. J Med Chem. 2021;64(10):6566–6568. doi:10.1021/acs.jmedchem.1c00698

20. Drost J, Clevers H. Organoids in cancer research. Nat Rev Cancer. 2018;18(7):407–418. doi:10.1038/s41568-018-0007-6

21. Ooft SN, Weeber F, Dijkstra KK, et al. Patient-derived organoids can predict response to chemotherapy in metastatic colorectal cancer patients. Sci Transl Med. 2019;11(513):eaay2574. doi:10.1126/scitranslmed.aay2574

22. Pasch CA, Favreau PF, Yueh AE, et al. Patient-Derived Cancer Organoid Cultures to Predict Sensitivity to Chemotherapy and Radiation. Clin Cancer Res. 2019;25(17):5376–5387. doi:10.1158/1078-0432.ccr-18-3590

23. Vlachogiannis G, Hedayat S, Vatsiou A, et al. Patient-derived organoids model treatment response of metastatic gastrointestinal cancers. Science. 2018;359(6378):920–926. doi:10.1126/science.aao2774

24. Lorenzi PL, Reinhold WC, Varma S, et al. DNA fingerprinting of the NCI-60 cell line panel. Mol Cancer Ther. 2009;8(4):713–724. doi:10.1158/1535-7163.mct-08-0921

25. Ianevski A, Giri AK, Aittokallio T. SynergyFinder 2.0: visual analytics of multi-drug combination synergies. Nucleic Acids Res. 2020;48(W1):W488–W493. doi:10.1093/nar/gkaa216

26. Bliss CI. THE TOXICITY OF POISONS APPLIED JOINTLY1. Ann Appl Biol. 1939;26(3):585–615. doi:10.1111/j.1744-7348.1939.tb06990.x

27. Yadav B, Wennerberg K, Aittokallio T, Tang J. Searching for Drug Synergy in Complex Dose–Response Landscapes Using an Interaction Potency Model. Comput Struct Biotechnology J. 2015;13:504–513. doi:10.1016/j.csbj.2015.09.001

28. Zhang DD, Hannink M. Distinct Cysteine Residues in Keap1 Are Required for Keap1-Dependent Ubiquitination of Nrf2 and for Stabilization of Nrf2 by Chemopreventive Agents and Oxidative Stress. Mol Cell Biol. 2003;23(22):8137–8151. doi:10.1128/mcb.23.22.8137-8151.2003

29. Kim MP, Evans DB, Wang H, Abbruzzese JL, Fleming JB, Gallick GE. Generation of orthotopic and heterotopic human pancreatic cancer xenografts in immunodeficient mice. Nat Protoc. 2009;4(11):1670–1680. doi:10.1038/nprot.2009.171

30. Roife D, Dai B, Kang Y, et al. Ex Vivo Testing of Patient-Derived Xenografts Mirrors the Clinical Outcome of Patients with Pancreatic Ductal Adenocarcinoma. Clin Cancer Res. 2016;22(24):6021–6030. doi:10.1158/1078-0432.ccr-15-2936

31. Evrard YA, Srivastava A, Randjelovic J, et al. Systematic Establishment of Robustness and Standards in Patient-Derived Xenograft Experiments and Analysis. Cancer Res. 2020;80(11):2286–2297. doi:10.1158/0008-5472.can-19-3101

32. van de Wetering M, Francies HE, Francis JM, et al. Prospective Derivation of a Living Organoid Biobank of Colorectal Cancer Patients. Cell. 2015;161(4):933–945. doi:10.1016/j.cell.2015.03.053

33. Miyoshi H, Stappenbeck TS. In vitro expansion and genetic modification of gastrointestinal stem cells in spheroid culture. Nat Protoc. 2013;8(12):2471–2482. doi:10.1038/nprot.2013.153

34. Klein M, Vignaud JM, Hennequin V, et al. Increased Expression of the Vascular Endothelial Growth Factor Is a Pejorative Prognosis Marker in Papillary Thyroid Carcinoma. J Clin Endocrinol Metabolism. 2001;86(2):656–658. doi:10.1210/jcem.86.2.7226

35. Bushnell B, Rood J, Singer E. BBMerge – Accurate paired shotgun read merging via overlap. Plos One. 2017;12(10):e0185056. doi:10.1371/journal.pone.0185056

36. Martin M. Cutadapt removes adapter sequences from high-throughput sequencing reads. Embnet J. 2011;17(1):10–12. doi:10.14806/ej.17.1.200

37. Dobin A, Davis CA, Schlesinger F, et al. STAR: ultrafast universal RNA-seq aligner. Bioinformatics. 2013;29(1):15–21. doi:10.1093/bioinformatics/bts635

38. Li B, Dewey CN. RSEM: accurate transcript quantification from RNA-Seq data with or without a reference genome. Bmc Bioinformatics. 2011;12(1):323. doi:10.1186/1471-2105-12-323

39. Love MI, Huber W, Anders S. Moderated estimation of fold change and dispersion for RNA-seq data with DESeq2. Genome Biol. 2014;15(12):550. doi:10.1186/s13059-014-0550-8

40. Subramanian A, Tamayo P, Mootha VK, et al. Gene set enrichment analysis: A knowledge-based approach for interpreting genome-wide expression profiles. Proc National Acad Sci. 2005;102(43):15545–15550. doi:10.1073/pnas.0506580102

41. Cerami E, Gao J, Dogrusoz U, et al. The cBio Cancer Genomics Portal: An Open Platform for Exploring Multidimensional Cancer Genomics Data. Cancer Discov. 2012;2(5):401–404. doi:10.1158/2159-8290.cd-12-0095

42. Gao J, Aksoy BA, Dogrusoz U, et al. Integrative Analysis of Complex Cancer Genomics and Clinical Profiles Using the cBioPortal. Sci Signal. 2013;6(269):pl1. doi:10.1126/scisignal.2004088

43. Giannakis M, Mu XJ, Shukla SA, et al. Genomic Correlates of Immune-Cell Infiltrates in Colorectal Carcinoma. Cell Reports. 2016;15(4):857–865. doi:10.1016/j.celrep.2016.03.075

44. Hoadley KA, Yau C, Hinoue T, et al. Cell-of-Origin Patterns Dominate the Molecular Classification of 10,000 Tumors from 33 Types of Cancer. Cell. 2018;173(2):291-304.e6. doi:10.1016/j.cell.2018.03.022

45. Vasaikar S, Huang C, Wang X, et al. Proteogenomic Analysis of Human Colon Cancer Reveals New Therapeutic Opportunities. Cell. 2019;177(4):1035-1049.e19. doi:10.1016/j.cell.2019.03.030

46. Tsherniak A, Vazquez F, Montgomery PG, et al. Defining a Cancer Dependency Map. Cell. 2017;170(3):564-576.e16. doi:10.1016/j.cell.2017.06.010

47. Gurusamy N, Rajasingh S, Sigamani V, et al. Noonan syndrome patient-specific induced cardiomyocyte model carrying SOS1 gene variant c.1654A>G. Exp Cell Res. 2021;400(1):112508. doi:10.1016/j.yexcr.2021.112508

48. Roberts AE, Araki T, Swanson KD, et al. Germline gain-of-function mutations in SOS1 cause Noonan syndrome. Nat Genet. 2007;39(1):70–74. doi:10.1038/ng1926

49. Tartaglia M, Pennacchio LA, Zhao C, et al. Gain-of-function SOS1 mutations cause a distinctive form of Noonan syndrome. Nat Genet. 2007;39(1):75–79. doi:10.1038/ng1939

50. Sanchez-Vega F, Mina M, Armenia J, et al. Oncogenic Signaling Pathways in The Cancer Genome Atlas. Cell. 2018;173(2):321-337.e10. doi:10.1016/j.cell.2018.03.035

51. Sheffels E, Sealover NE, Wang C, et al. Oncogenic RAS isoforms show a hierarchical requirement for the guanine nucleotide exchange factor SOS2 to mediate cell transformation. Sci Signal. 2018;11(546):eaar8371. doi:10.1126/scisignal.aar8371

52. Nichols RJ, Haderk F, Stahlhut C, et al. RAS nucleotide cycling underlies the SHP2 phosphatase dependence of mutant BRAF-, NF1- and RAS-driven cancers. Nat Cell Biol. 2018;20(9):1064–1073. doi:10.1038/s41556-018-0169-1

53. Kuzu OF, Noory MA, Robertson GP. The Role of Cholesterol in Cancer. Cancer Res. 2016;76(8):2063–2070. doi:10.1158/0008-5472.can-15-2613

54. Solanki HS, Welsh EA, Fang B, et al. Cell Type–specific Adaptive Signaling Responses to KRASG12C Inhibition. Clin Cancer Res. Published online 2021. doi:10.1158/1078-0432.ccr-20-3872

55. Soleimani A, Rahmani F, Ferns GA, Ryzhikov M, Avan A, Hassanian SM. Role of the NF-κB signaling pathway in the pathogenesis of colorectal cancer. Gene. 2019;726:144132. doi:10.1016/j.gene.2019.144132

