## Supplemental Figures for "The translational role of SOS1 in colorectal cancer"

**Supplemental Table 1.** Baseline characteristics of patients whose tumor samples were used for CRC PDO development

| PDO model | Age | Race | Gender | Tumor location | Stage | KRAS | MSI-H/MMR deficient |
|---|---|---|---|---|---|---|---|
| MCC19990-002 | 69 | black | female | colon | III | G12A | no |
| MCC19990-004 | 56 | white | male | colon | II | A146T | yes |
| MCC19990-006 | 58 | hispanic | male | colon | I | G12C | no |
| MCC19990-007 | 78 | white | female | colon | II | wild type | yes |
| MCC19990-010 | 75 | white | female | colon | II | G12A | no |
| MCC19990-013 | 59 | hispanic | female | colon | III | G12A | no |
| MCC20584-004 | 73 | white | female | peritoneum | IV | G12S | no |
| MCC20584-005 | 48 | white | female | Abdominal wall | IV | wild type | no |
| MCC20584-010 | 65 | white | male | liver | IV | G12D | yes |

Note: tumors with available genomic information were *BRAF* wild-type and *HER2* non-amplified.

**Supplemental Figure 1.** Molecular aberrations, mRNA, and protein expressions of SOS1 stratified by clinical and histopathologic factors in CRC. **(A)** Prevalence of *SOS1* mutations across different stages of CRC in DFCI CRC cohort is not statistically different. $p$=0.2. **(B)** The frequency of *SOS1* mutations compared to that of common genes in primary and metastatic CRC in GENIE cohort v11.0. **(C)** Prevalence of *SOS1* mutations in right versus left CRC in DFCI CRC cohort is not statistically different. $p$=0.3. **(D)** The frequency of *SOS1* alterations compared to that of *KRAS, TP53, BRAF, APC, SMAD4, PI3KCA* and *HER2* alterations in CRC of GENIE cohort v11.0. *SOS1* has co-occurrence with these genes in CRC with p-value <0.001 and q-value <0.001 (Benjamini-Hochberg FDR correction). Of note, unprofiled samples for any of these genes were excluded. Samples with no alterations in any of these genes were excluded from the figure. **(E)** *SOS1* and *SOS2* mutually co-occurrence in DFCI CRC cohort. *SOS1* has co-occurrence with *SOS2* in CRC with p-value = 0.034 and q-value =0.034. **(F)** *SOS1* mRNA expression (RSEM, Batch normalized from Illumina HiSeq_RNASeqV2, $log_2$) in *RAS/RAF* wild-type versus *RAS/RAF* mutant CRC in TCGA PanCancer Atlas. mRNA expression as $log_2$(value + 1), $p$=0.3. **(G)** *SOS1* mRNA expression (RSEM, Batch normalized from Illumina HiSeq_RNASeqV2, $log_2$) across different stages of CRC in TCGA PanCancer Atlas. mRNA expression as $log_2$(value + 1), $p$=1. **(H)** SOS1 protein expression (mass spectrometry by CPTAC) in *RAS/RAF* wild-type versus *RAS/RAF* mutant CRC in CPTAC-2 Prospective. $p$=0.3. **(I)** SOS1 protein expression (mass spectrometry by CPTAC) across different stages of CRC in CPTAC-2 Prospective. $p$=0.4. **(J)** Correlation between SOS1 and SOS2 protein expression (mass spectrometry by CPTAC) in CRC in CPTAC-2 Prospective. Spearman's ρ: 0.45, $p$=0.1.

**A.**

| Tumor stage | SOS1 mutant | SOS1 wild-type | *p* value |
|---|---|---|---|
| I | 9 (5.9%) | 143 (94.1%) | 0.2 |
| II | 9 (4.8%) | 178 (95.2%) | |
| III | 3 (1.9%) | 156 (98.1%) | |
| IV | 1 (1.5%) | 64 (98.5%) | |

**B.**

| Molecular alterations | Primary CRC | Metastatic CRC | *p value* |
|---|---|---|---|
| SOS1 (mutant/wild-type) | 3.2% (126/3870) | 2.3% (38/1634) | 0.07 |
| KRAS (mutant/wild-type) | 43.6% (3620/4684) | 43.9% (1868/2383) | 0.7 |
| BRAF (mutant/wild-type) | 12% (995/7307) | 8.2% (348/3901) | <0.001 |
| HER2 (amplified/wild-type) | 2.4% (123/5051) | 2.6% (65/2474) | 0.6 |

**C.**

| Tumor sites | SOS1 mutant | SOS1 wild-type | *p* value |
|---|---|---|---|
| Right | 14 (4.4%) | 301 (95.6%) | 0.3 |
| Left | 8 (2.6%) | 295 (97.4%) | |

**D.**

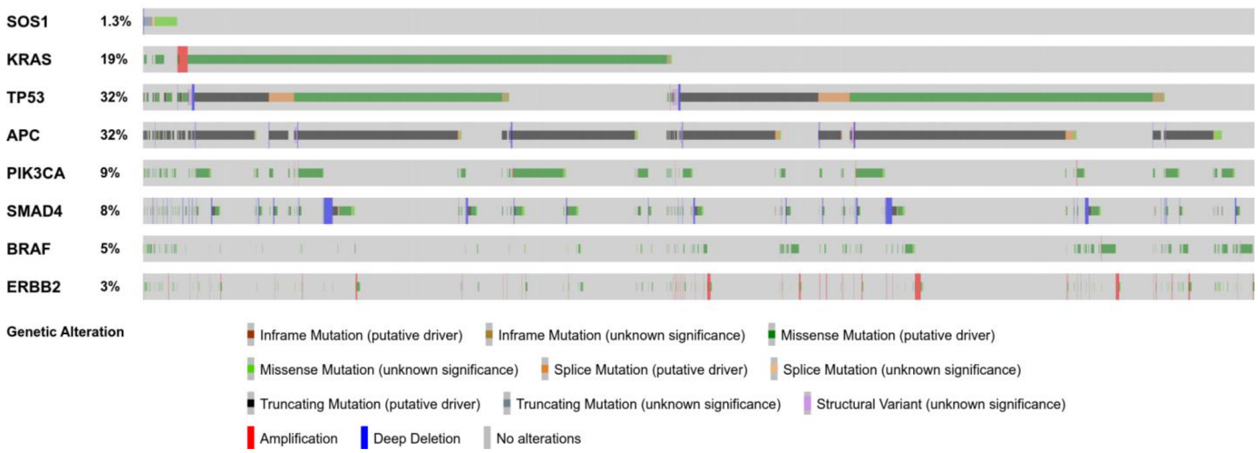


**E.**

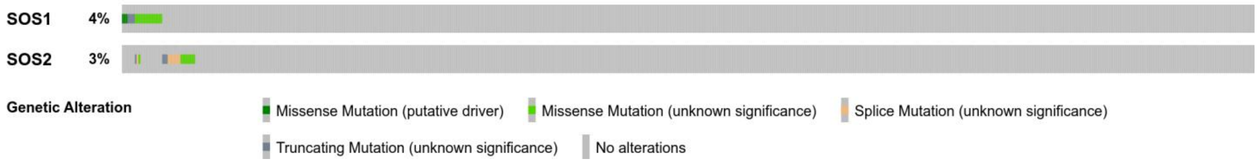

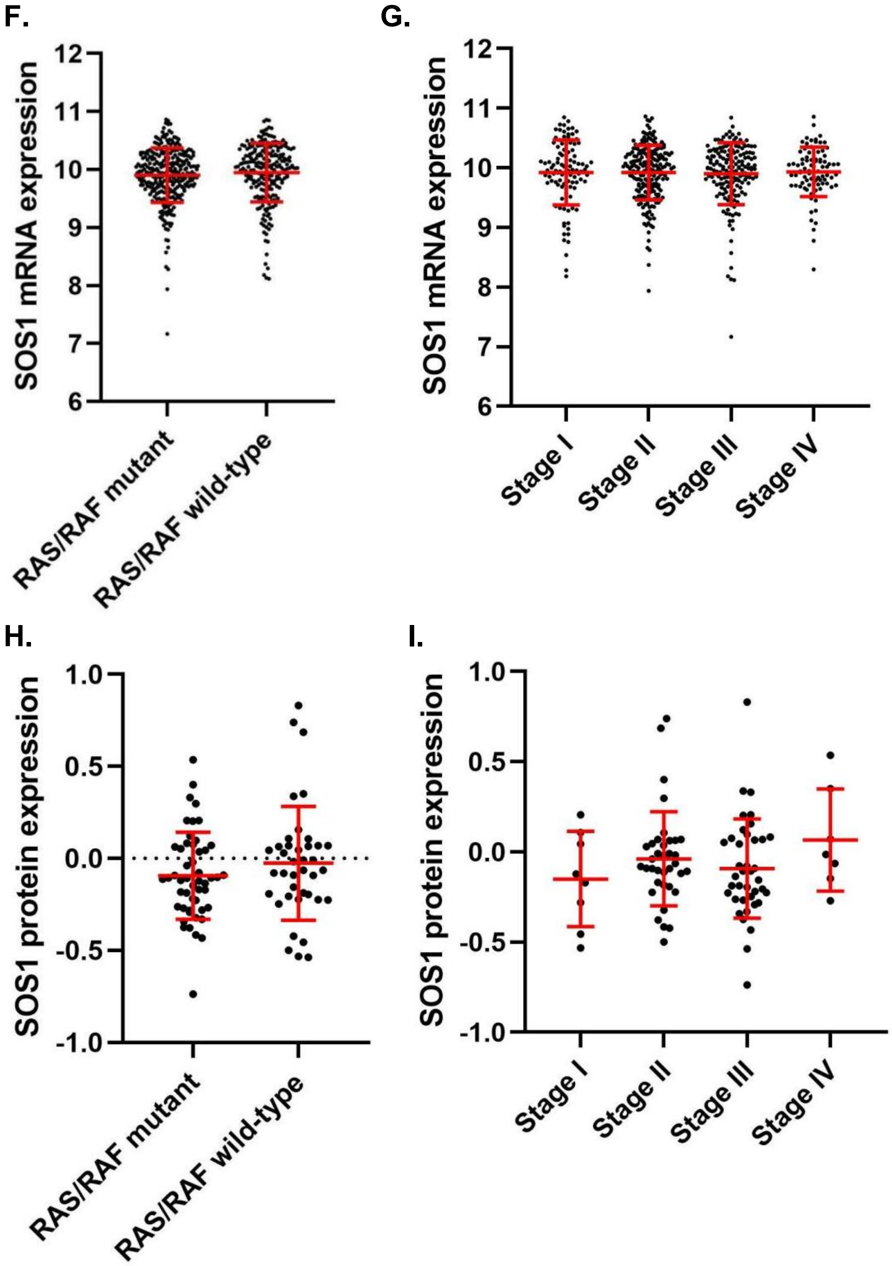
F.
G.
SOS1 mRNA expression
12
11
10
9
8
7
6
RAS/RAF mutant
RAS/RAF wild-type
Stage I
Stage II
Stage III
Stage IV
H.
I.
SOS1 protein expression
1.0
0.5
0.0
-0.5
-1.0
RAS/RAF mutant
RAS/RAF wild-type
Stage I
Stage II
Stage III
Stage IV

**J.**

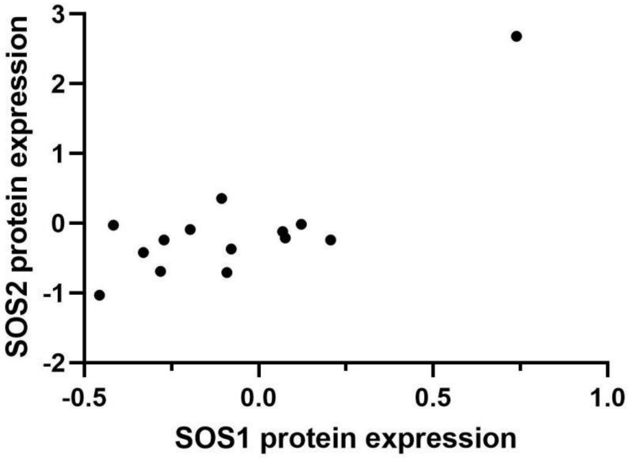


**Supplemental Figure 2.** SOS1 and SOS2 expressions by IHC in CRC PDX models. **(A)** SOS1 expression by IHC across different PDX generations 1-3. *p*=0.5. **(B)** SOS2 expression by IHC across different PDX generations 1-3. *p*=0.7. **(C)** SOS1/SOS2 expression ratio by IHC across different PDX generations 1-3. *p*=0.5. **(D)** SOS1 expression by IHC between primary and metastatic CRC PDX models. *p*=0.8. **(E)** SOS2 expression by IHC between primary and metastatic CRC PDX models. *p*=0.4. **(F)** SOS1/SOS2 expression ratio by IHC between primary and metastatic CRC PDX models. *p*=0.4. **(G)** SOS1 expression by IHC between *KRAS* wild-type and *KRAS* mutant CRC PDX models. *p*=1. **(H)** SOS2 expression by IHC between *KRAS* wild-type and *KRAS* mutant CRC PDX models. *p*=0.5. **(I)** SOS1/SOS2 expression ratio by IHC between *KRAS* wild-type and *KRAS* mutant CRC PDX models. *p*=0.4. **(J)** CRC PDX tumor growth measured as adjusted AUC in association with SOS1 expression. *p*=0.3. **(K)** CRC PDX tumor growth measured as adjusted AUC in association with SOS2 expression. SOS2 H-score group (≥6 vs <6): *p*=1.0. **(L)** CRC PDX tumor growth measured as adjusted AUC in association with SOS1/SOS2 expression ratio. SOS1/SOS2 ratio group (>1 vs ≤1): *p*=1.0.

**A.**

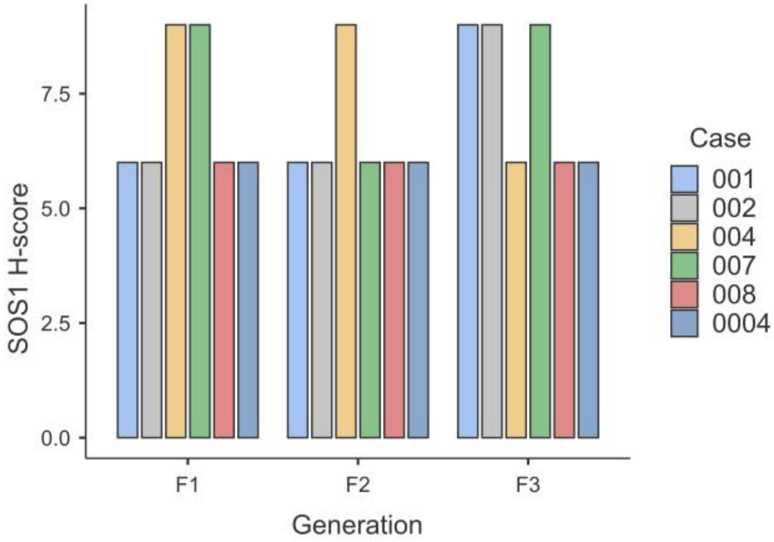

**B.**

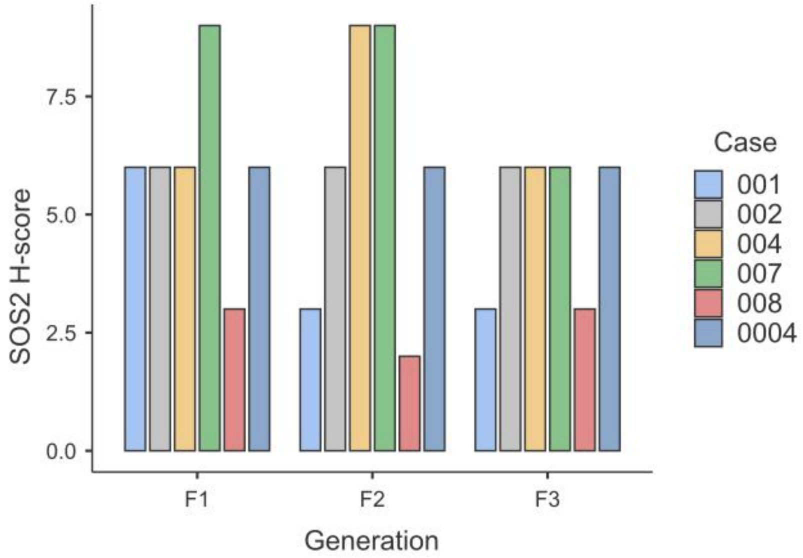
7.5
5.0
2.5
0.0
SOS2 H-score
F1
F2
F3
Generation
Case
001
002
004
007
008
0004

**C.**

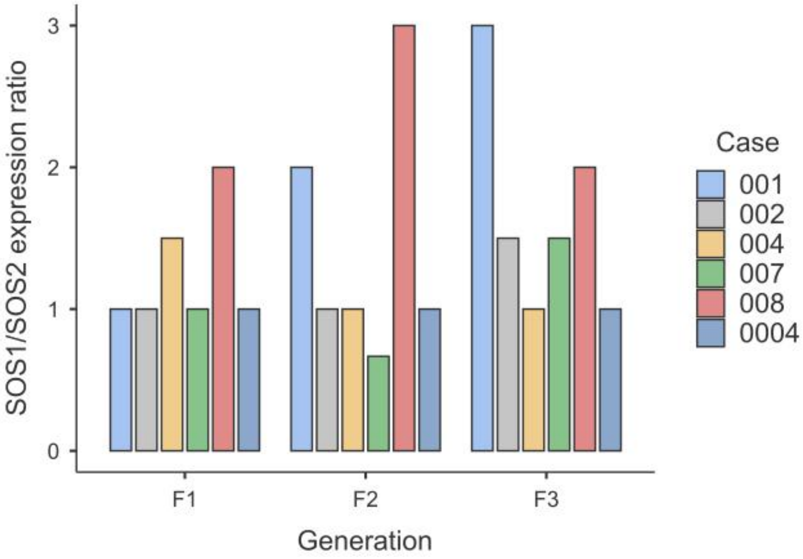
3
2
1
0
SOS1/SOS2 expression ratio
F1
F2
F3
Generation
Case
001
002
004
007
008
0004

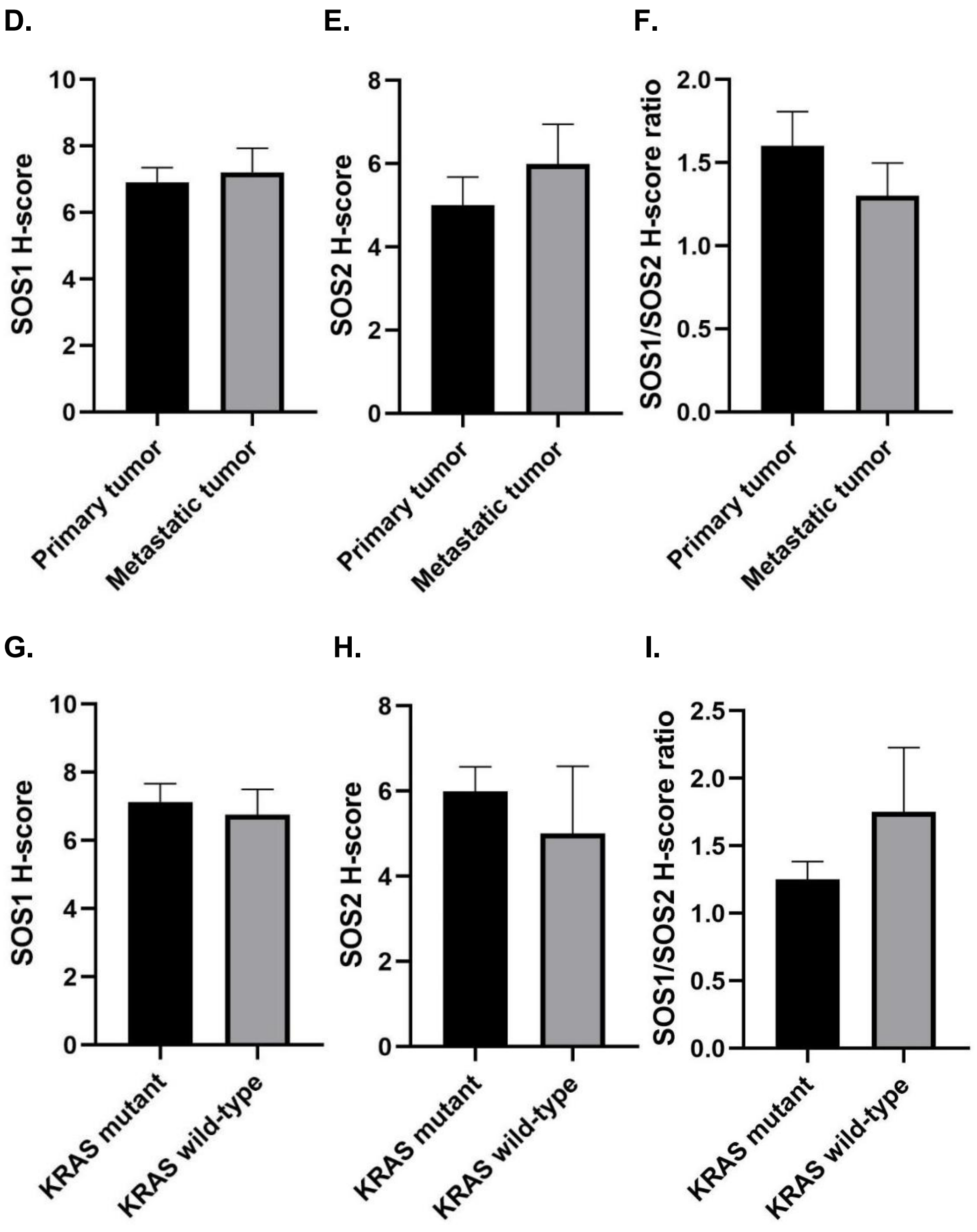
D.
E.
F.
SOS1 H-score
SOS2 H-score
SOS1/SOS2 H-score ratio
Primary tumor
Metastatic tumor
G.
H.
I.
KRAS mutant
KRAS wild-type
0
2
4
6
8
10
0.0
0.5
1.0
1.5
2.0
2.5

J. K. L.

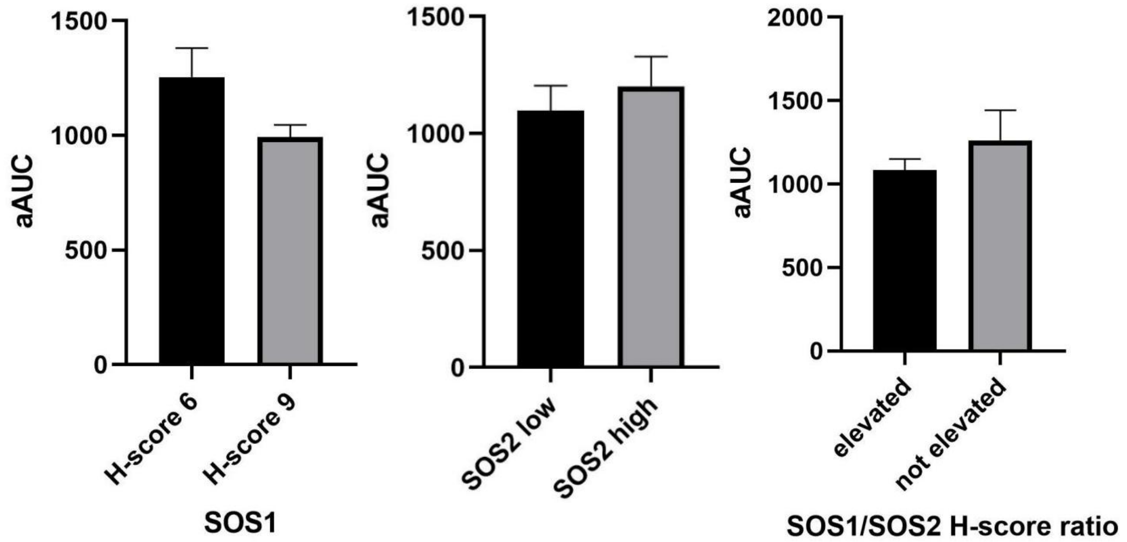


**Supplemental Figure 3.** mRNA expression and gene set enrichment analyses of CRC cell lines and CRC PDOs. **(A)** Gene expressions of CRC cell lines with differential sensitivity to SOS1 inhibitor BI3406. The sensitivity was defined by Hoffmann et al, Cancer Discovery, 2021. The dataset has 23088 features (genes). # of markers for phenotype sensitive: 11075 (48.0%) with correlation area 48.0%. # of markers for phenotype resistant: 12013 (52.0%) with correlation area 52.0%. Heatmap of the top 50 features for each phenotype sensitive (n=3) or resistant (n=5) to BI3406. **(B)** Enrichment plots in BI3406 sensitive CRC cell lines (n=3) vs resistant CRC cell lines (n=5). 30/50 gene sets are upregulated in phenotype "sensitive". 9 gene sets are significant at FDR < 25%. 8 gene sets are significantly enriched at nominal p value < 1%. 9 gene sets are significantly enriched at nominal p value < 5%. 20/50 gene sets are upregulated in phenotype "resistant". 1 gene sets are significantly enriched at FDR < 25%. 1 gene sets are significantly enriched at nominal p value < 1%. 1 gene sets are significantly enriched at nominal p value < 5%. **(C)** Gene expressions of CRC cell lines with or without SOS1 dependency. The dataset has 19177 features (genes). # of markers for phenotype 1:7177 (37.4%) with correlation area 32.6%. # of markers for phenotype 0: 12000 (62.6% ) with correlation area 67.4%. Heatmap of the top 50 features for each phenotype dependent (n=8) or independent (n=40) to *SOS1*. **(D)** Enrichment plots in SOS1 dependent CRC cell lines (n=8) vs independent CRC cell lines (n=40). 30/50 gene sets are upregulated in phenotype SOS1 dependent. 23 gene sets are significant at FDR < 25%. 17 gene sets are significantly enriched at nominal p value < 1%. 19 gene sets are significantly enriched at nominal p value < 5%. 20/50 gene sets are upregulated in phenotype SOS1 independent. 1 gene sets are significantly enriched at FDR < 25%. 0 gene sets are significantly enriched at nominal p value < 1%. 1 gene sets are significantly enriched at nominal p value < 5%. **(E)** GSEA of CRC cell lines with differential sensitivity to SOS1 inhibition, *SOS1* dependency, and *KRAS* dependency.

A.

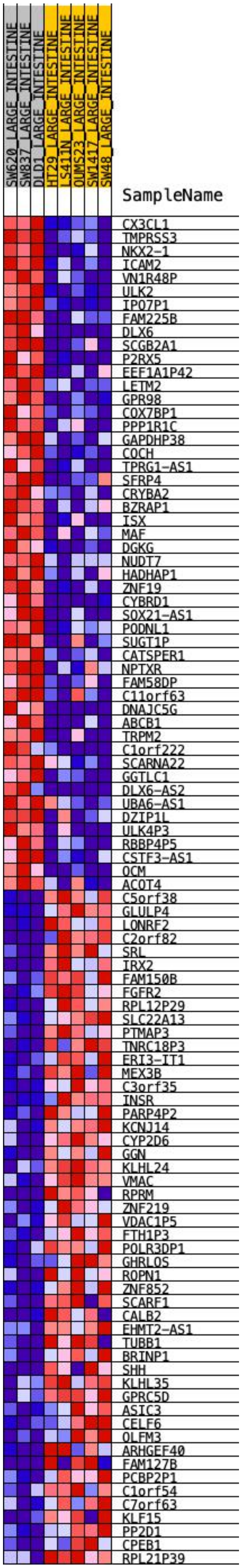
SW620_LARGE_INTESTINE
SW837_LARGE_INTESTINE
DLD1_LARGE_INTESTINE
HT29_LARGE_INTESTINE
LS411N_LARGE_INTESTINE
OUMS23_LARGE_INTESTINE
SW1417_LARGE_INTESTINE
SW48_LARGE_INTESTINE
SampleName
CX3CL1
TMPRSS3
NKX2-1
ICAM2
VN1R48P
ULK2
IPO7P1
FAM225B
DLX6
SCGB2A1
P2RX5
EEF1A1P42
LETM2
GPR98
COX7BP1
PPP1R1C
GAPDHP38
COCH
TPRG1-AS1
SFRP4
CRYBA2
BZRAP1
ISX
MAF
DGKG
NUDT7
HADHAP1
ZNF19
CYBRD1
SOX21-AS1
PODNL1
SUGT1P
CATSPER1
NPTXR
FAM58DP
C11orf63
DNAJC5G
ABCB1
TRPM2
C1orf222
SCARNA22
GGTLC1
DLX6-AS2
UBA6-AS1
DZIP1L
ULK4P3
RBBP4P5
CSTF3-AS1
OCM
ACOT4
C5orf38
GLULP4
LONRF2
C2orf82
SRL
IRX2
FAM150B
FGFR2
RPL12P29
SLC22A13
PTMAP3
TNRC18P3
ERI3-IT1
MEX3B
C3orf35
INSR
PARP4P2
KCNJ14
CYP2D6
GGN
KLHL24
VMAC
RPRM
ZNF219
VDAC1P5
FTH1P3
POLR3DP1
GHRLOS
ROPN1
ZNF852
SCARF1
CALB2
EHMT2-AS1
TUBB1
BRINP1
SHH
KLHL35
GPRC5D
ASIC3
CELF6
OLFM3
ARHGEF40
FAM127B
PCBP2P1
C1orf54
C7orf63
KLF15
PP2D1
CPEB1
RPL21P39

**B.**

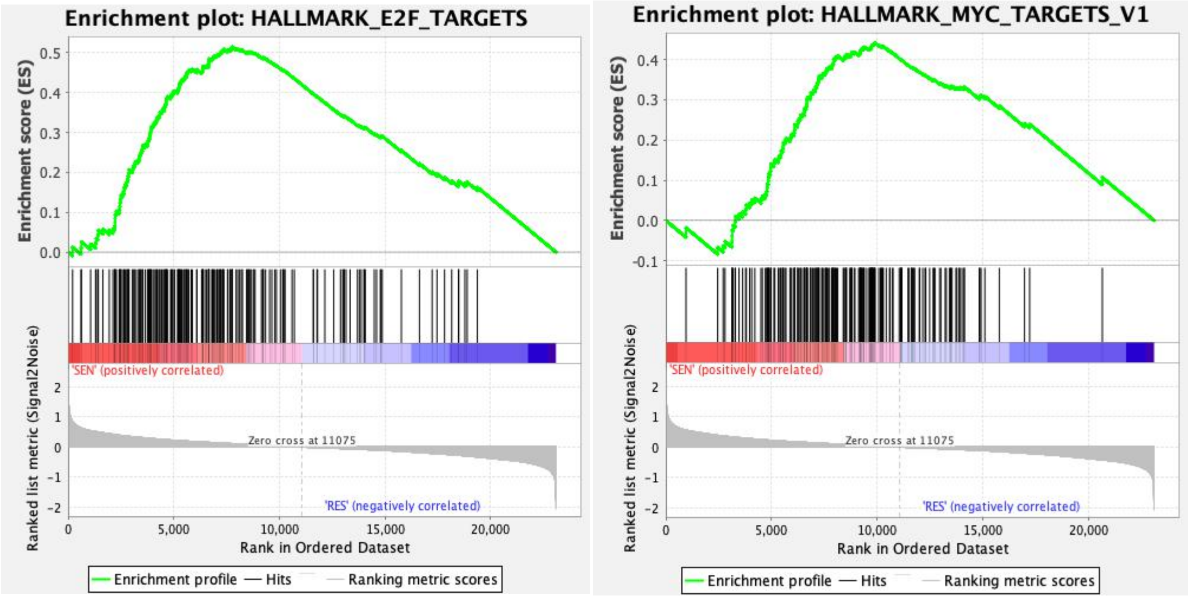

Enrichment plot: HALLMARK_E2F_TARGETS
Enrichment score (ES)
0.5
0.4
0.3
0.2
0.1
0.0
'SEN' (positively correlated)
Ranked list metric (Signal2Noise)
2
1
0
-1
-2
Zero cross at 11075
'RES' (negatively correlated)
0
5,000
10,000
15,000
20,000
Rank in Ordered Dataset
Enrichment profile
Hits
Ranking metric scores
Enrichment plot: HALLMARK_MYC_TARGETS_V1
Enrichment score (ES)
0.4
0.3
0.2
0.1
0.0
-0.1
'SEN' (positively correlated)
Ranked list metric (Signal2Noise)
2
1
0
-1
-2
Zero cross at 11075
'RES' (negatively correlated)
0
5,000
10,000
15,000
20,000
Rank in Ordered Dataset
Enrichment profile
Hits
Ranking metric scores

**C.**

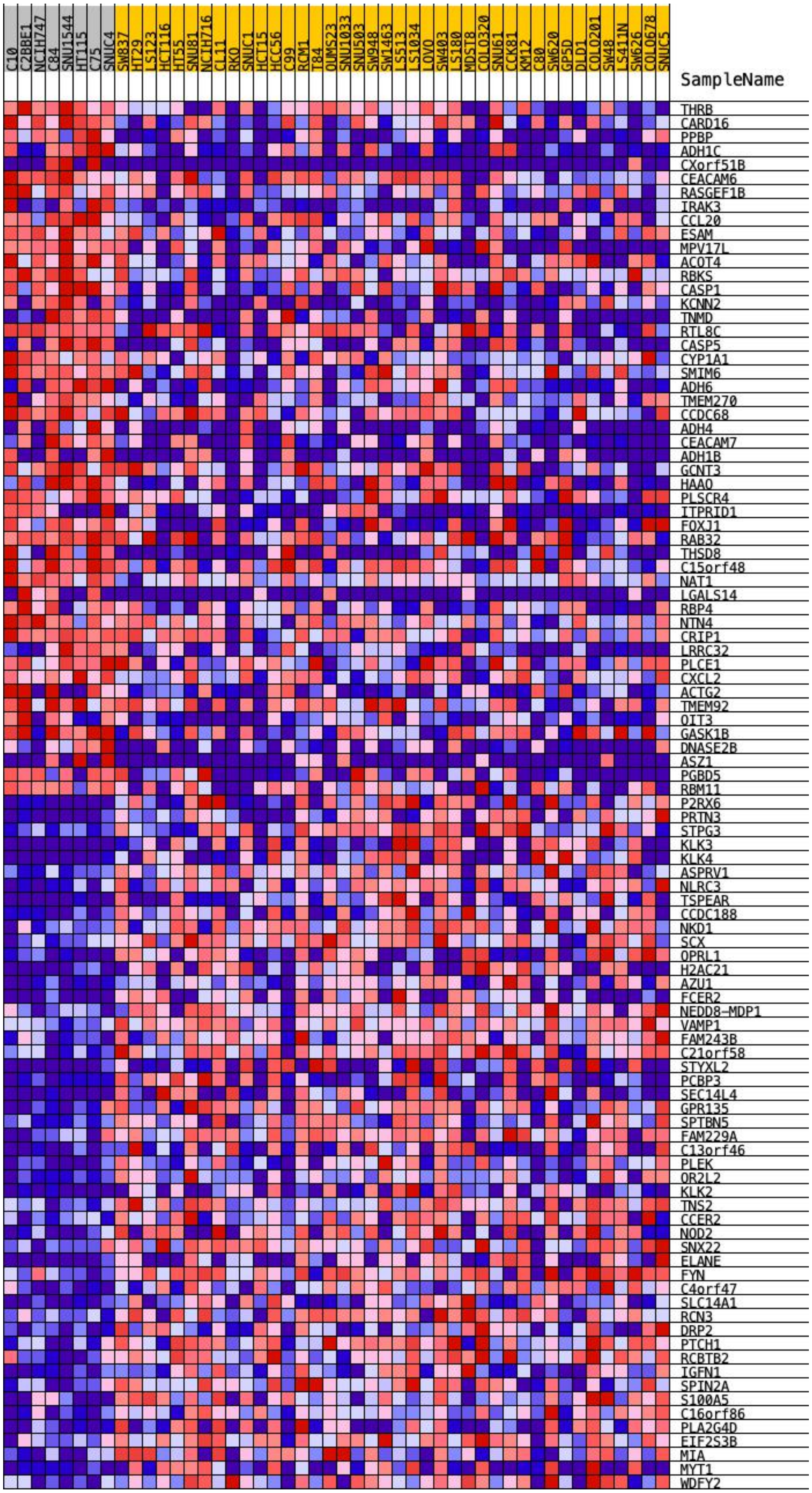
C10
C2BBE1
NCIH747
C84
SNU1544
HT115
C75
SNUC4
SW837
HT29
LS123
HCT116
HT55
SNU81
NCIH716
CL11
RKO
SNUC1
HCT15
HCC56
C99
RCM1
T84
OUMS23
SNU1033
SNU503
SW948
SW1463
LS513
LS1034
LOVO
SW403
LS180
MDST8
COLO320
SNU61
CCK81
KM12
C80
SW620
GP5D
DLD1
COLO201
SW48
LS411N
SW626
COLO678
SNUC5
SampleName
THRB
CARD16
PPBP
ADH1C
CXorf51B
CEACAM6
RASGEF1B
IRAK3
CCL20
ESAM
MPV17L
ACOT4
RBKS
CASP1
KCNN2
TNMD
RTL8C
CASP5
CYP1A1
SMIM6
ADH6
TMEM270
CCDC68
ADH4
CEACAM7
ADH1B
GCNT3
HAAO
PLSCR4
ITPRID1
FOXJ1
RAB32
THSD8
C15orf48
NAT1
LGALS14
RBP4
NTN4
CRIP1
LRRC32
PLCE1
CXCL2
ACTG2
TMEM92
OIT3
GASK1B
DNASE2B
ASZ1
PGBD5
RBM11
P2RX6
PRTN3
STPG3
KLK3
KLK4
ASPRV1
NLRC3
TSPEAR
CCDC188
NKD1
SCX
OPRL1
H2AC21
AZU1
FCER2
NEDD8-MDP1
VAMP1
FAM243B
C21orf58
STYXL2
PCBP3
SEC14L4
GPR135
SPTBN5
FAM229A
C13orf46
PLEK
OR2L2
KLK2
TNS2
CCER2
NOD2
SNX22
ELANE
FYN
C4orf47
SLC14A1
RCN3
DRP2
PTCH1
RCBTB2
IGFN1
SPIN2A
S100A5
C16orf86
PLA2G4D
EIF2S3B
MIA
MYT1
WDFY2

**D.**

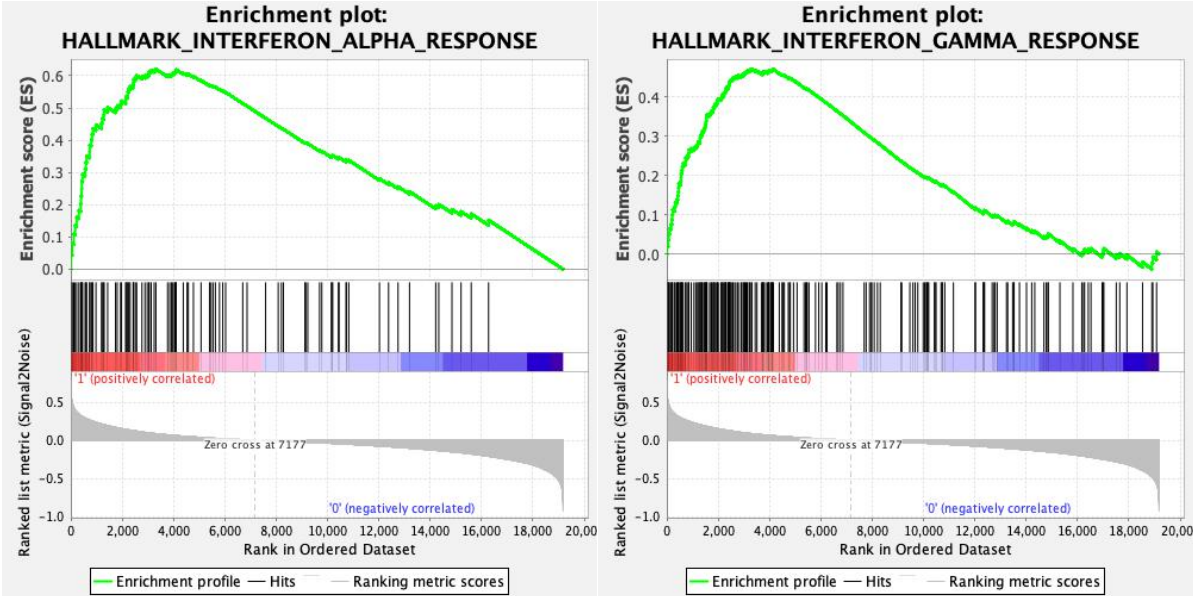


**E.**

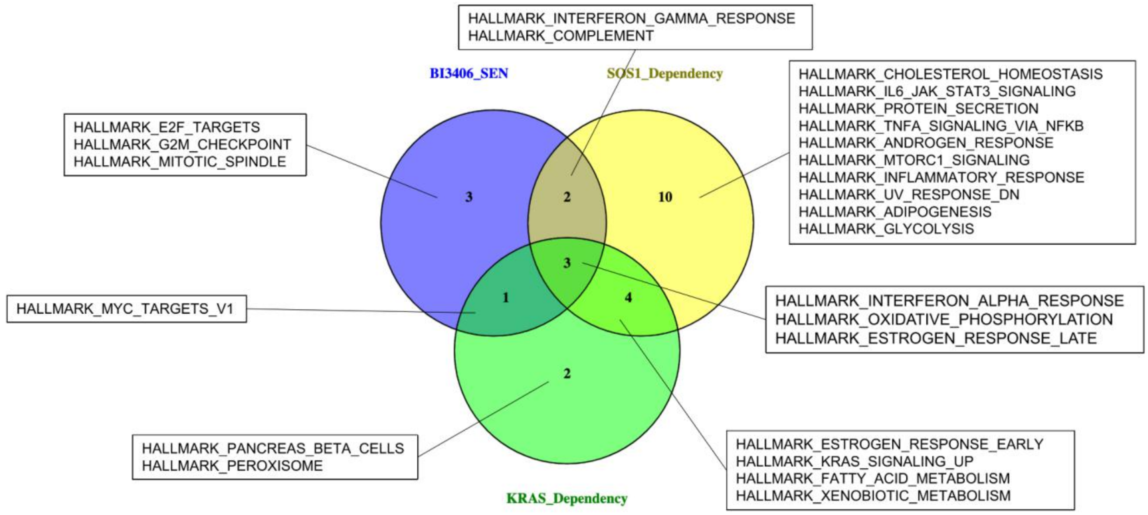


**Supplemental Figure 4.** Markers not predictive to *SOS1* dependency or sensitivity to SOS1 inhibitor BI3406 in CRC models. **(A)** Association of SOS1 protein expression with *SOS1* dependency in DepMap database. Spearman's ρ -0.069, $p$=0.75. **(B)** Association of *SOS1* mRNA expression ($\log_2$(TPM+1), Expression 21Q4 Public) with *SOS1* dependency in DepMap database. Spearman's ρ -0.013, $p$=0.9. **(C)** Association of *SOS1/SOS2* mRNA expression ratio with *SOS1* dependency in DepMap database. Spearman's ρ -0.156, $p$=0.14. Note: $H_a$ is a negative correlation. **(D)** SOS1 expression by IHC between BI3406 sensitive and BI3406 resistant CRC PDX. $p$=1. **(E)** SOS2 expression by IHC between BI3406 sensitive and BI3406 resistant CRC PDX. $p$=0.3.

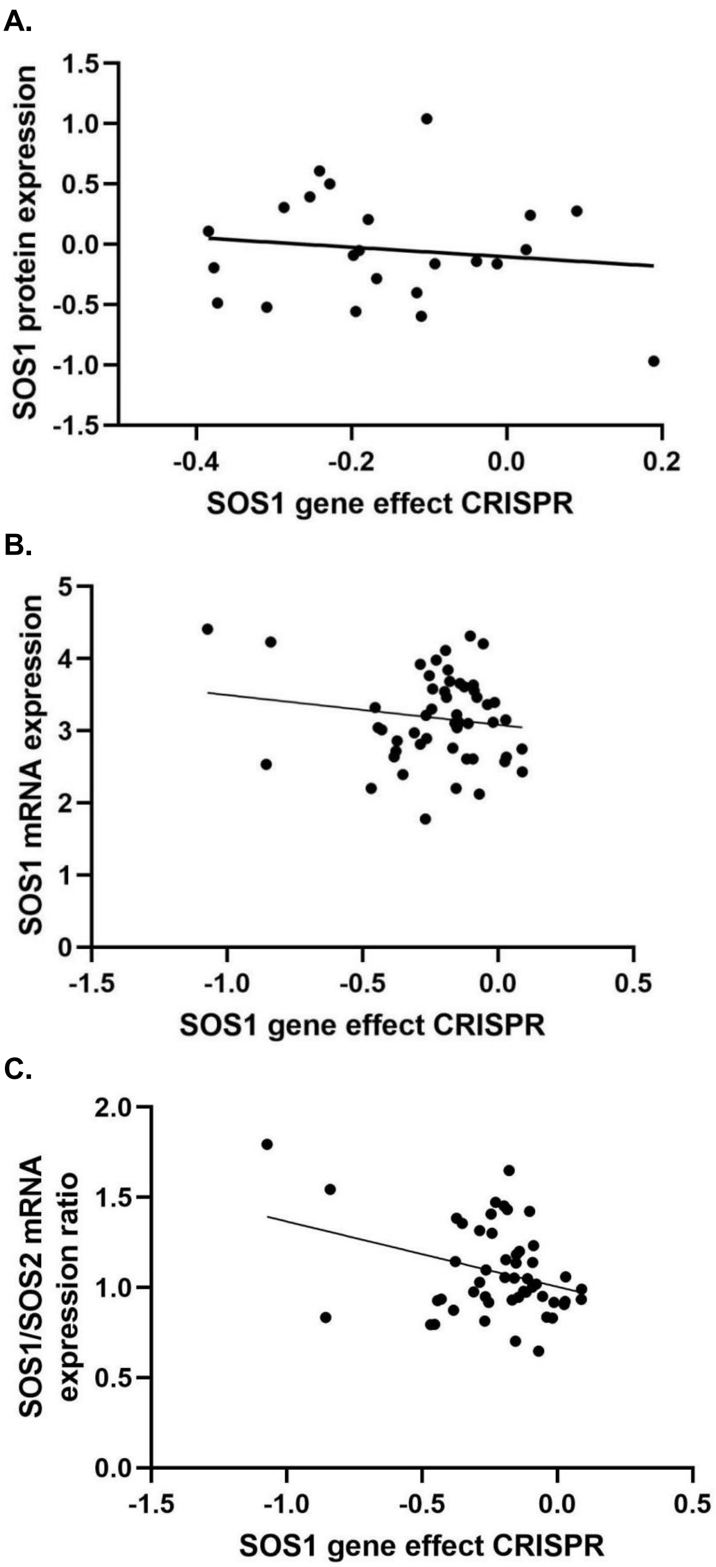
A.
SOS1 protein expression
1.5
1.0
0.5
0.0
-0.5
-1.0
-1.5
-0.4
-0.2
0.0
0.2
SOS1 gene effect CRISPR
B.
SOS1 mRNA expression
5
4
3
2
1
0
-1.5
-1.0
-0.5
0.0
0.5
SOS1 gene effect CRISPR
C.
SOS1/SOS2 mRNA expression ratio
2.0
1.5
1.0
0.5
0.0
-1.5
-1.0
-0.5
0.0
0.5
SOS1 gene effect CRISPR

**D.**

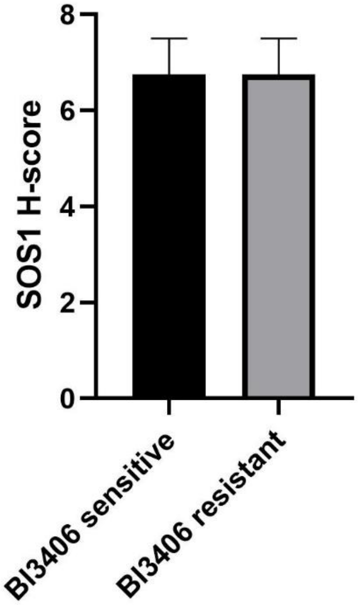


**E.**

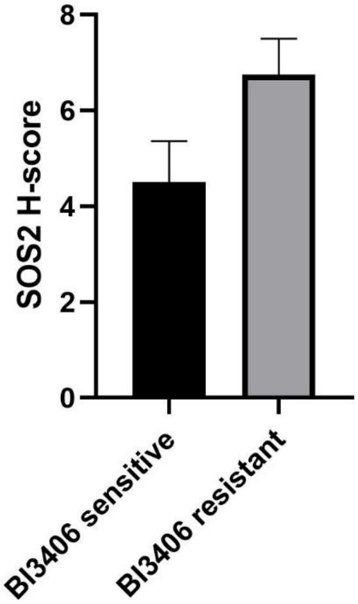


**Supplemental Figure 5.** Effect of SOS1 inhibitor BI3406 on CRC cells and PDOs. **(A)** SOS1 and pERK levels of HCT116 after treatment with BI3406 at various concentrations for 6 hours or 24 hours. **(B)** Summary of GSEA of BI3406-sensitive CRC PDOs before and after treatment with 1µM BI3406 for 24 hours. **(C)** *SOS1* knockdown with siRNA after 48 hours was associated with increased apoptosis in SOS1 inhibitor BI3406 resistant CRC cell line HCT116. **(D)** Relative viability of HCT116 after treatment with various concentrations of buparlisib in the presence or absence of BI3406 for 72 hours. BI3406 has no synergistic cytotoxic effect with buparlisib. **(E)** Relative viability of HCT116 after treatment with various concentrations of defactinib in the presence or absence of BI3406 for 72 hours. BI3406 has no synergistic cytotoxic effect with defactinib.

**A.**

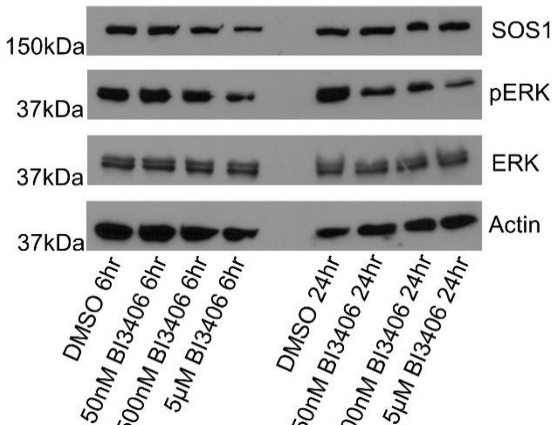

**B.**

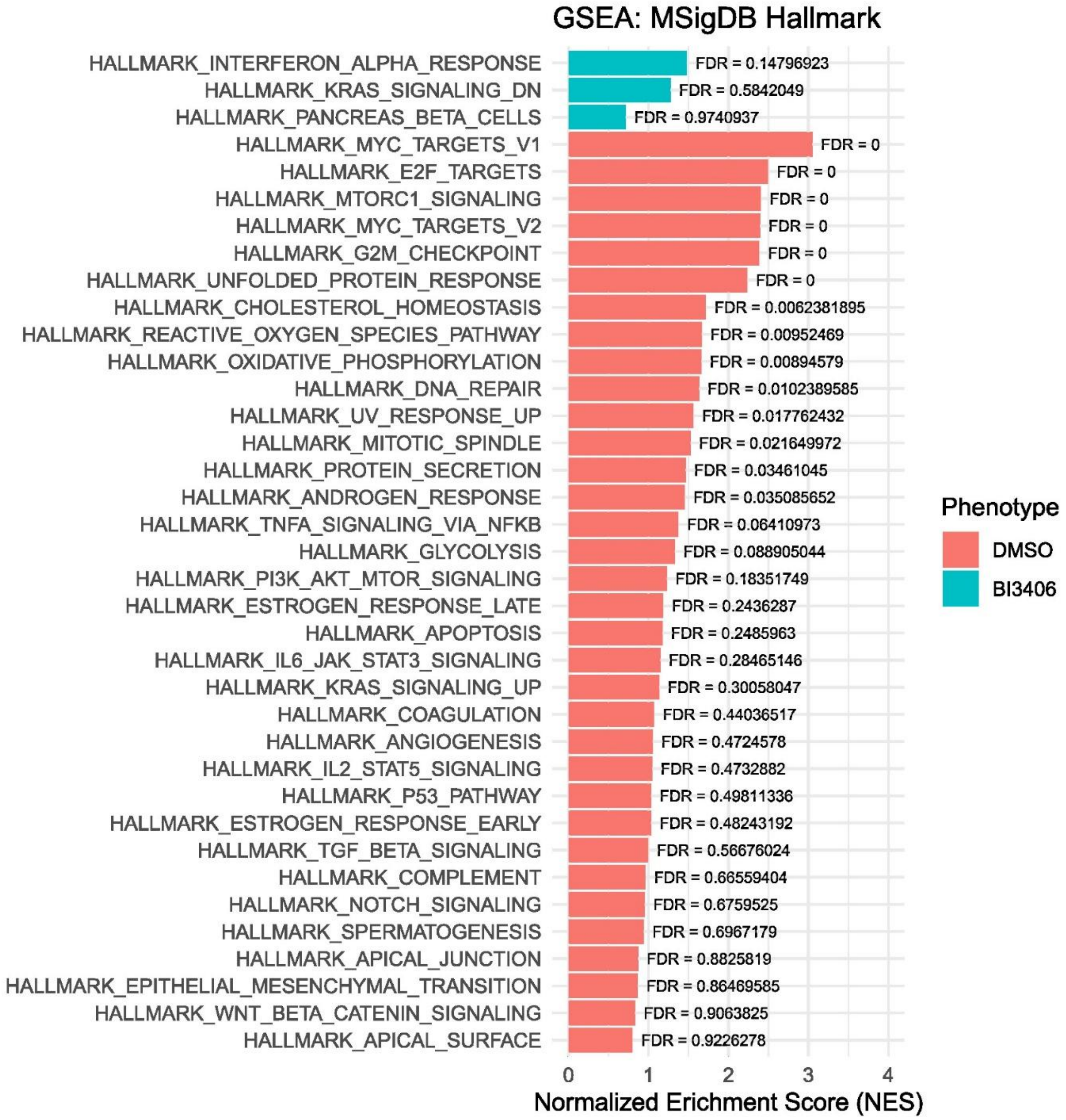

GSEA: MSigDB Hallmark
HALLMARK_INTERFERON_ALPHA_RESPONSE
FDR = 0.14796923
HALLMARK_KRAS_SIGNALING_DN
FDR = 0.5842049
HALLMARK_PANCREAS_BETA_CELLS
FDR = 0.9740937
HALLMARK_MYC_TARGETS_V1
FDR = 0
HALLMARK_E2F_TARGETS
FDR = 0
HALLMARK_MTORC1_SIGNALING
FDR = 0
HALLMARK_MYC_TARGETS_V2
FDR = 0
HALLMARK_G2M_CHECKPOINT
FDR = 0
HALLMARK_UNFOLDED_PROTEIN_RESPONSE
FDR = 0
HALLMARK_CHOLESTEROL_HOMEOSTASIS
FDR = 0.0062381895
HALLMARK_REACTIVE_OXYGEN_SPECIES_PATHWAY
FDR = 0.00952469
HALLMARK_OXIDATIVE_PHOSPHORYLATION
FDR = 0.00894579
HALLMARK_DNA_REPAIR
FDR = 0.0102389585
HALLMARK_UV_RESPONSE_UP
FDR = 0.017762432
HALLMARK_MITOTIC_SPINDLE
FDR = 0.021649972
HALLMARK_PROTEIN_SECRETION
FDR = 0.03461045
HALLMARK_ANDROGEN_RESPONSE
FDR = 0.035085652
HALLMARK_TNFA_SIGNALING_VIA_NFKB
FDR = 0.06410973
HALLMARK_GLYCOLYSIS
FDR = 0.088905044
HALLMARK_PI3K_AKT_MTOR_SIGNALING
FDR = 0.18351749
HALLMARK_ESTROGEN_RESPONSE_LATE
FDR = 0.2436287
HALLMARK_APOPTOSIS
FDR = 0.2485963
HALLMARK_IL6_JAK_STAT3_SIGNALING
FDR = 0.28465146
HALLMARK_KRAS_SIGNALING_UP
FDR = 0.30058047
HALLMARK_COAGULATION
FDR = 0.44036517
HALLMARK_ANGIOGENESIS
FDR = 0.4724578
HALLMARK_IL2_STAT5_SIGNALING
FDR = 0.4732882
HALLMARK_P53_PATHWAY
FDR = 0.49811336
HALLMARK_ESTROGEN_RESPONSE_EARLY
FDR = 0.48243192
HALLMARK_TGF_BETA_SIGNALING
FDR = 0.56676024
HALLMARK_COMPLEMENT
FDR = 0.66559404
HALLMARK_NOTCH_SIGNALING
FDR = 0.6759525
HALLMARK_SPERMATOGENESIS
FDR = 0.6967179
HALLMARK_APICAL_JUNCTION
FDR = 0.8825819
HALLMARK_EPITHELIAL_MESENCHYMAL_TRANSITION
FDR = 0.86469585
HALLMARK_WNT_BETA_CATENIN_SIGNALING
FDR = 0.9063825
HALLMARK_APICAL_SURFACE
FDR = 0.9226278
0
1
2
3
4
Normalized Erichment Score (NES)
Phenotype
DMSO
BI3406

C.

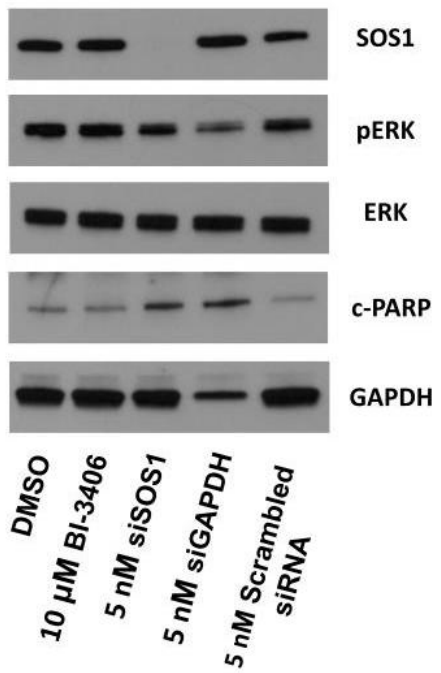
SOS1
pERK
ERK
c-PARP
GAPDH
DMSO
10 μM BI-3406
5 nM siSOS1
5 nM siGAPDH
5 nM Scrambled siRNA

D.

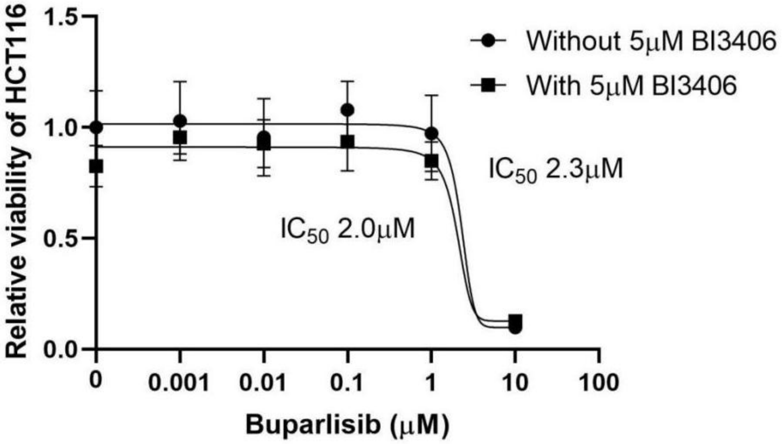
Without 5μM BI3406
With 5μM BI3406
IC50 2.3μM
IC50 2.0μM
Relative viability of HCT116
1.5
1.0
0.5
0.0
0
0.001
0.01
0.1
1
10
100
Buparlisib (μM)

E.

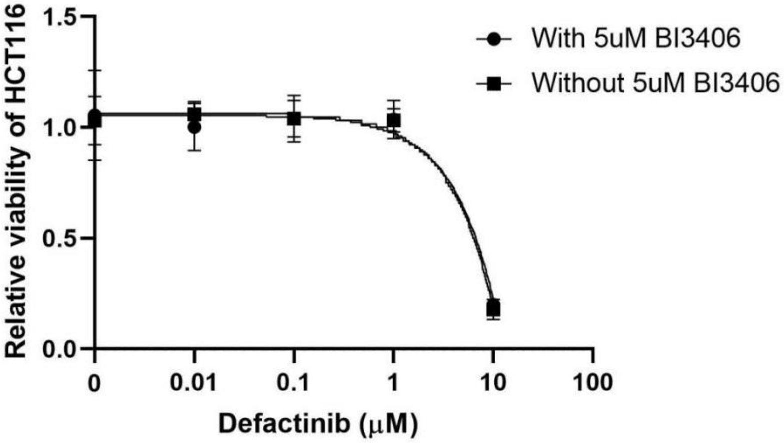

Relative viability of HCT116
1.5
1.0
0.5
0.0
0
0.01
0.1
1
10
100
Defactinib (μM)
With 5uM BI3406
Without 5uM BI3406
